# Time evolution of PEG-shedding and serum protein coronation determines the cell uptake kinetics and delivery of lipid nanoparticle formulated mRNA

**DOI:** 10.1101/2021.08.20.457104

**Authors:** A. Gallud, M. J. Munson, K. Liu, A. Idström, H. M. G. Barriga, S. R. Tabaei, N. Aliakbarinodehi, M. Ojansivu, Q. Lubart, J. J. Doutch, M. N. Holme, L. Evenäs, L. Lindfors, M. M. Stevens, A. Collén, A. Sabirsh, F. Höök, E. K. Esbjörner

**Affiliations:** Division of Chemical Biology, Department of Biology and Biological Engineering, Chalmers University of Technology, Gothenburg, Sweden; Advanced Drug Delivery, Pharmaceutical Sciences, R&D, AstraZeneca, Gothenburg, Sweden; Department of Chemistry and Chemical Engineering, Chalmers University of Technology, Gothenburg, Sweden; Department of Medical Biochemistry and Biophysics, Karolinska Institutet, SE-171 77 Stockholm, Sweden; Department of Physics, Chalmers University of Technology, 41296 Gothenburg, Sweden; ISIS Neutron and Muon Source, STFC, Rutherford Appleton Laboratory, Didcot OX11 ODE, U.K.; Imperial College London, Materials, London, SW7 2AZ. U.K.; Projects, Research and Early Development, Cardiovascular, Renal and Metabolism (CVRM), Biopharmaceuticals R&D, AstraZeneca, Gothenburg, Sweden

**Keywords:** Lipid nanoparticle, mRNA delivery, serum, PEG-shedding, protein corona, uptake kinetics, pulse-chase imaging, apolipoprotein

## Abstract

Development of efficient lipid nanoparticle (LNP) vectors remains a major challenge towards broad clinical translation of RNA therapeutics. New lipids will be required, but also better understanding LNP interactions with the biological environment. Herein, we model protein corona formation on PEG-ylated DLin-MC3-DMA LNPs and identify time-dependent maturation steps that critically unlock their cellular uptake and mRNA delivery. Uptake requires active serum proteins and precedes after a significant (∼2 hours) lag-time, which we show can be eliminated by pre-incubating LNPs for 3-4 hours in serum-containing media. This indicates an important role of protein corona maturation for the pharmacokinetic effects of these LNPs. We show, using single-nanoparticle imaging, NMR diffusometry, SANS, and proteomics, that the LNPs, upon serum exposure, undergo rapid PEG-shedding (∼30 minutes), followed by a slower rearrangement of the adsorbed protein layer. The PEG-shedding coincides in time with high surface abundance of Apolipoprotein A-II, whereas the LNPs preferentially bind Apolipoprotein E when their maximum uptake-competent state is reached. Finally, we show that pre-incubation of the LNPs enables rapid uptake and allows pulse-chase video-microscopy colocalization experiments with sufficiently short pulse durations to gain improved mechanistic understanding of how intracellular trafficking events determine delivery efficacy, emphasizing early endosomes as important delivery-mediating compartments.

## 1. Introduction

RNA-based therapeutics are gaining significant momentum due to their ability to specifically target protein expression *in vivo*, thereby opening up new possibilities to treat currently undruggable diseases in multiple areas.[1, 2] mRNA drugs are particularly attractive as they enable high levels of *in situ* protein production, thereby circumventing current manufacturing and stability drawbacks associated with protein replacement therapeutics; their significant potential has also been recently highlighted by the approval of mRNA vaccines against Covid- 19 during the ongoing SARS-Cov2-2 pandemic.[3] The major challenge towards translating additional new RNA modalities into clinical use, particularly in the protein replacement therapy area, lies in the development of methods offering efficient delivery of these high molecular weight and heavily charged therapeutics into target cells. Recent developments in nanotechnology and material science have yielded many promising non-viral delivery systems for safe and non-invasive RNA delivery.[4] Lipid nanoparticle (LNP) based vehicles have so far been most successful[5] underlying for example the first FDA approved siRNA therapeutic product, Onpattro, used for the treatment of hereditary transthyretin amyloidosis,[6–8] and the Comirnaty and mRNA-1273 mRNA vaccines from respectively Pfizer/BioNTech[9] and Moderna[10] which are now used to combat the Covid-19 pandemic. Still, significant challenges remain to enable broadly applicable delivery solutions for mRNA therapeutics, particularly in situations that requires long-term repeated intravenous administration and therefore places extra demands on safety and delivery efficacy.

LNPs can be easily formulated to efficiently encapsulate both siRNA and mRNA using microfluidic mixing[11–14] and thereby protect the RNA from nucleases, promote delivery across the cell membrane and facilitate endosomal escape.[5, 15–18] LNPs are typically comprised of an ionizable cationic lipid to encapsulate RNA and aid intracellular delivery with low toxicity,[19, 20] combined with several “helper lipids”,[21] such as phospholipids, a sterol, and a lipidated polyethylene glycol (PEG; PEG-ylated lipid). In addition to the introduction of new ionizable lipids,[17, 22–27] LNP efficacy can be significantly modulated by fine-tuning of LNP lipid compositions to induce changes of physicochemical parameters such as size and surface properties[28] or by chemical modification of individual helper components such as cholesterol.[29] Optimisation can result in both improved total cell uptake and, importantly, better endosomal escape.[30] This is, to some extent, further complicated by the fact that LNPs, just like other engineered nanomaterials,[31–34] are interacting with their biological environment prior to and during uptake,[35] leading to both the formation of functionally important protein coronas and modifications to the LNP particles themselves, [36, 37] for example via shedding (release) of the PEG-ylated lipid, herein referred to as PEG-shedding.

PEG-ylated phospholipids are commonly incorporated into lipid-based drug carriers as they offer a “stealth” effect to increase the circulation time of the drug product in the body. The PEG-ylated phospholipids can also be designed to regulate LNP size and transfection efficiency.[28, 38] For example, the acyl chain length of the PEG-lipid, its degree of saturation, and concentration can influence both *in vitro* and *in vivo* functional response of siRNA-loaded LNPs,[39, 40] presumably because the lipid tail properties alter the propensity of PEG-lipids to shed from the surface of LNPs upon interaction with serum proteins, as observed by pulsed gradient spin echo NMR,[41] and hence loss of steric shielding.

As mentioned above, *in vitro* and *in vivo* nanoparticle delivery efficiency is often greatly influenced by the formation of a protein adsorption layer (commonly referred to as a protein corona; although lipids and glycans may associate as well).[42, 43] Once formed, the protein corona contributes to define the nanoparticle’s surface and hence interaction potential with the biological environment.[35] The protein adsorption layer can be divided into a “hard” corona defined by high affinity and low exchange rate and a “soft” corona with lower affinity and higher exchange rate. It has been suggested that the protein corona plays an important role in nanoparticle biodistribution, cellular uptake, and toxicity[44] and it is therefore crucial to consider it alongside “native” properties of LNPs. In a limited number of recent studies, the protein coronas of LNPs have been investigated and the efficiency of LNPs containing DLin- MC3-DMA as the ionizable lipid component is reported to be dependent on the coronal accumulation of apolipoprotein E (ApoE).[36, 37, 45] Interestingly, LNPs containing DLin-MC3- DMA with a modified backbone structure (i.e. alkyne and ester groups introduced into the lipid tails), led to an ApoE-independent LNP cellular uptake pathway in the liver, in favor of albumin-associated macropinocytosis and endocytosis.[23] This certainly demonstrates the complexity of, and need to investigate, the corona. It also opens the possibility to use the corona as a way to target LNPs to cells or tissues.[46] Importantly, the characterization of protein corona formation commonly rely on samples from single time points and do not explore the evolution of the corona with time. Furthermore, no study has so far addressed both PEG-shedding and protein corona formation on the LNP surface to pinpoint how the dynamics of these two processes jointly modulate cell uptake and subsequent cargo delivery.

In the present work, we have studied how mRNA-encapsulating LNPs, containing DLin-MC3- DMA and DMPE-PEG2000 to provide steric shielding, interact with serum proteins prior to cell uptake *in vitro*. Our study particularly focuses on modelling the temporal development of a diverse protein corona on the surface of LNPs and on exploring how this affects cell uptake kinetics and delivery efficacy. We show that the cell uptake of the LNPs is strongly serum-dependent and proceeds first after a significant lag time (∼2 hours) during which protein coronation occurs. Using a combination of different biophysical assays to monitor protein coronation and PEG-shedding, we show that the LNPs, when exposed to serum, undergo rapid PEG-shedding (concomitant with the build-up of a first protein corona) and thereafter a significantly slower protein corona rearrangement step. In parallel, we develop a new protocol to pre-treat LNPs with serum-containing media prior to cell exposure. With this we demonstrate that significant time (3-4 hours) is needed for the LNPs to adopt their most uptake-competent state pointing out the importance of protein corona maturation to unlock cell entry and subsequently mRNA-delivery efficiency of these LNPs. Finally, we demonstrate that pre-incubation of LNPs in serum enables rapid uptake during very short cell exposures. This allows pulse-chase time-lapse experiments with sufficient time resolution to study the intracellular trafficking of LNPs; the latter being a significant factor to generate the fundamental knowledge required to succeed in the development of next generation nucleotide-based drug delivery.

## 2. Results and discussion

### 2.1. Pre-incubation of LNPs in serum-containing culture media enhances the uptake kinetics and onset of protein expression following mRNA delivery

To explore the role of serum proteins in the cellular uptake and cytosolic delivery of PEG- ylated and DLin-MC3-DMA lipid containing mRNA-loaded lipid nanoparticles (LNPs) were used. These were composed of DLin-MC3-DMA, DSPC, cholesterol, and DMPE_PEG at molar ratios of 50:10:38.5:1.5 (**supporting Table S1**) and hereafter referred to as MC3_DMPE_PEG. The encapsulated mRNA cargo encoded for eGFP; a mixture of Cy5- labelled mRNA (20%) unlabelled mRNA (80%) to ensure both good visualization and efficient protein translation. Prior to cell experiments, the LNPs were characterised with respect to size (85±5 nm in diameter) and monodispersity (PDI<0.05) using dynamic light scattering. The RNA encapsulation, measured using the RiboGreen™ RNA assay, was determined to 98% (**supporting Table S1, SI text**).

The LNPs were diluted in cell culture media with different concentrations (0-10%) of fetal bovine serum (FBS) and added to cultures of human Huh-7 hepatic cells for time-lapse monitoring of mRNA uptake and translation (**Movie S1**). **Figure 1a** shows representative overlay images of the Cy5 and eGFP signals for different FBS concentrations after 7 hours of incubation. At this time point both Cy5 uptake and eGFP expression was clearly visible if serum proteins were present. It has been shown that Huh-7 cells can secrete apolipoprotein E (ApoE), which has been pointed out as important for MC3-DLin-DMA LNP uptake via the LDLR receptor.[45] Consistent with this, we also found some LNP uptake and eGFP expression after 17 hours (at the end of the video recordings; **Movie S1**) in the serum free cell cultures. Nevertheless, presence of serum proteins in the culture media resulted in significantly faster uptake kinetics (**Movie S1**) and the higher eGFP levels clearly persist also at the end-point (**Figure S2**). This demonstrates that the presence of serum proteins is important for productive delivery of mRNA mediated by MC3-containing LNPs.

**Figure 1.**
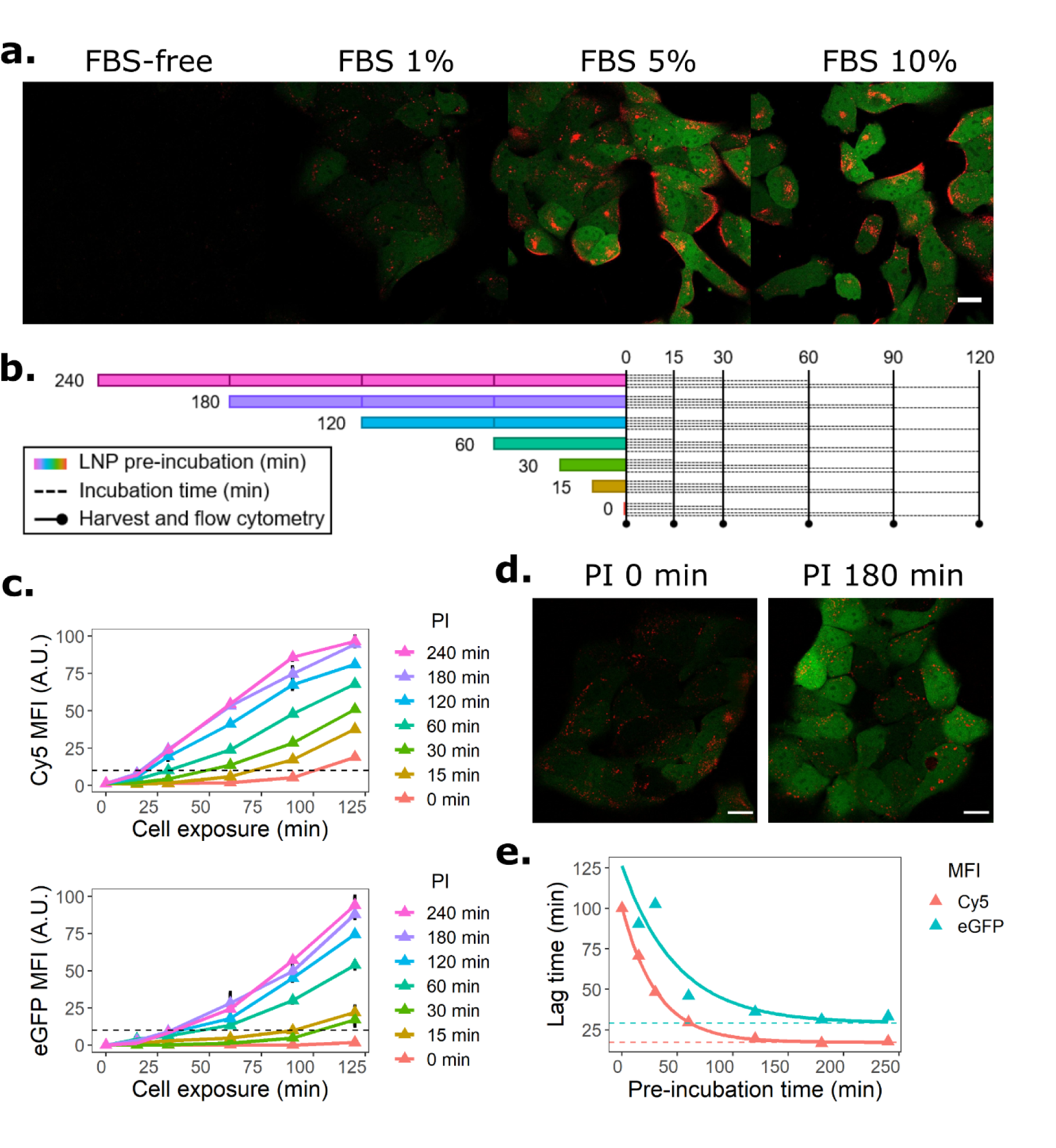
Pre-incubation of LNPs in serum-containing media accelerates mRNA uptake kinetics and onset of protein production. **(a)** Confocal microscopy images of Huh-7 cells showing the cellular uptake (red; Cy5-labelled mRNA) and protein expression (green; eGFP protein) following continuous exposure to LNPs for 7 hours in presence of different concentrations of fetal bovine serum (FBS). **(b)** Schematic of the experimental setup to evaluate the effect of LNP pre-incubation (PI) on the kinetics of LNP cellular uptake and subsequent protein expression. **(c)** Cellular uptake of Cy5-labelled mRNA (top) and corresponding cellular eGFP fluorescence (bottom) as function of cell exposure time, following increasing durations of pre-incubation at 37°C (PI, time indicated in legend) of the LNPs in cell culture media with 10% FBS; the data were collected according to the schematic description in (b). The cellular Cy5 and eGFP fluorescence intensities were quantified by flow cytometry on live cells (error bars are standard deviations of the sampling distribution from triplicate performed in three independent experiments). The dashed lines represent 10% of the maximum Cy5/eGFP signal, see further in (e). **(d)** Confocal microscopy images of cells exposed to fresh (not pre-incubated; PI 0 min) or 180 minutes pre-incubated (PI 180 min) LNPs (red: Cy5; green: eGFP). **(e)** Lag time in LNP uptake (Cy5 signal) and protein expression (eGFP signal) as function of pre-incubation (PI) time. The lag time was defined as the time taken to reach 10% of the maximum MFI signal in (b). Single exponential decay functions were fitted to the data (solid lines). MC3_DMPE_PEG LNPs at a concentration of 0.625 µg/mL (mRNA basis) were used for all experiments. The scale bars in all images are 20 µm.

These observations inspired us to pre-incubate LNPs in FBS-containing cell culture medium for 0 to 240 minutes to allow serum protein interactions with the LNPs to establish prior to cell exposure (**Figure 1b**). The pre-incubated LNPs were added to Huh-7 cells and the intracellular Cy5 and eGFP signals were quantified at discrete time points for up to 125 minutes post-exposure (**Figure 1c-d; Supporting Figure S3**). The data show that both uptake kinetics and evolution of the eGFP expression is strongly dependent on the pre-incubation (PI) time. In absence of pre-incubation (0 minute data), the uptake, and consequently protein production, has a clear lag time, indicating that the LNPs must interact with serum components to adopt an uptake-competent state and that this process takes significant time (∼2 hours). Figure 1e shows that the lag time of detectable cell uptake (Cy5) and protein expression (eGFP) decays exponentially with pre-incubation time, reaching minimum plateau values of 17 and 29 minutes for Cy5 and eGFP respectively. This suggests that the time delay between cell uptake and functional delivery is short, and would be consistent with a delivery mechanism by which endosomal escape and cytosolic release occurs rapidly following cell entry, as proposed by Wittrup et al. for the cellular delivery of siRNA.[47, 48]

It has indeed been suggested that PEG-lipid functionalised LNPs must shed at least parts of the PEG layer prior to cell entry;[41] which is consistent with the herein observation of a lag-time in cellular uptake. In addition, upon interaction with serum, LNPs will adopt a diverse and LNP- composition-sensitive protein corona.[35] Our finding that LNPs need to be pre-incubated for a significant period of time (240 minutes) to reach their optimal uptake-competent state, indicates that the temporal evolution of these two processes, possibly also involving a configurational/compositional rearrangement of the corona[32, 35] is likely to take place. Interestingly, extending the pre-incubation time to 24 hours did not increase the LNP uptake compared to 240 minutes, and actually halved the production of eGFP (**Figure S4**), suggesting that an optimum exists. As detailed in the forthcoming sections, the likely interplay between protein-corona formation and PEG-shedding, and the reduced eGFP production at prolonged serum incubation times prompted us to undertake further biophysical and cell-based investigations of the serum-mediated effect on LNP uptake.

### 2.2. Effect of temperature and serum heat-inactivation on delivery efficiency of pre-incubated LNPs

To address configurational and conformational aspects of the protein corona formed on the surface of LNPs, we repeated the experiment outlined in Figure 1b using high content imaging-based screening.[30] This enabled a combinatorial investigation of the effect of pre-incubation time (0-3 hours), pre-incubation temperatures (22-42℃), and normal versus heat-inactivated FBS (**Figure 2**). We also compared the response of the LNPs with DMPE_PEG lipid (C14 acyl chain) to LNPs with DSPE_PEG lipid (C18 acyl chain; **supporting Table S1**), both having saturated acyl chains. The latter screen showed that LNP containing DSPE_PEG was completely unable to enter Huh-7 hepatic cells regardless of the pre-incubation conditions, and therefore did not lead to any eGFP expression (**Figure S5**). This is consistent with the suggestion that C18 saturated acyl chains mediate firmer attachment of the PEG-lipid to the LNP[39] and strongly suggests that PEG-shedding (i.e. the release of PEG lipids from the LNPs) must occur prior to cell uptake.

**Figure 2.**
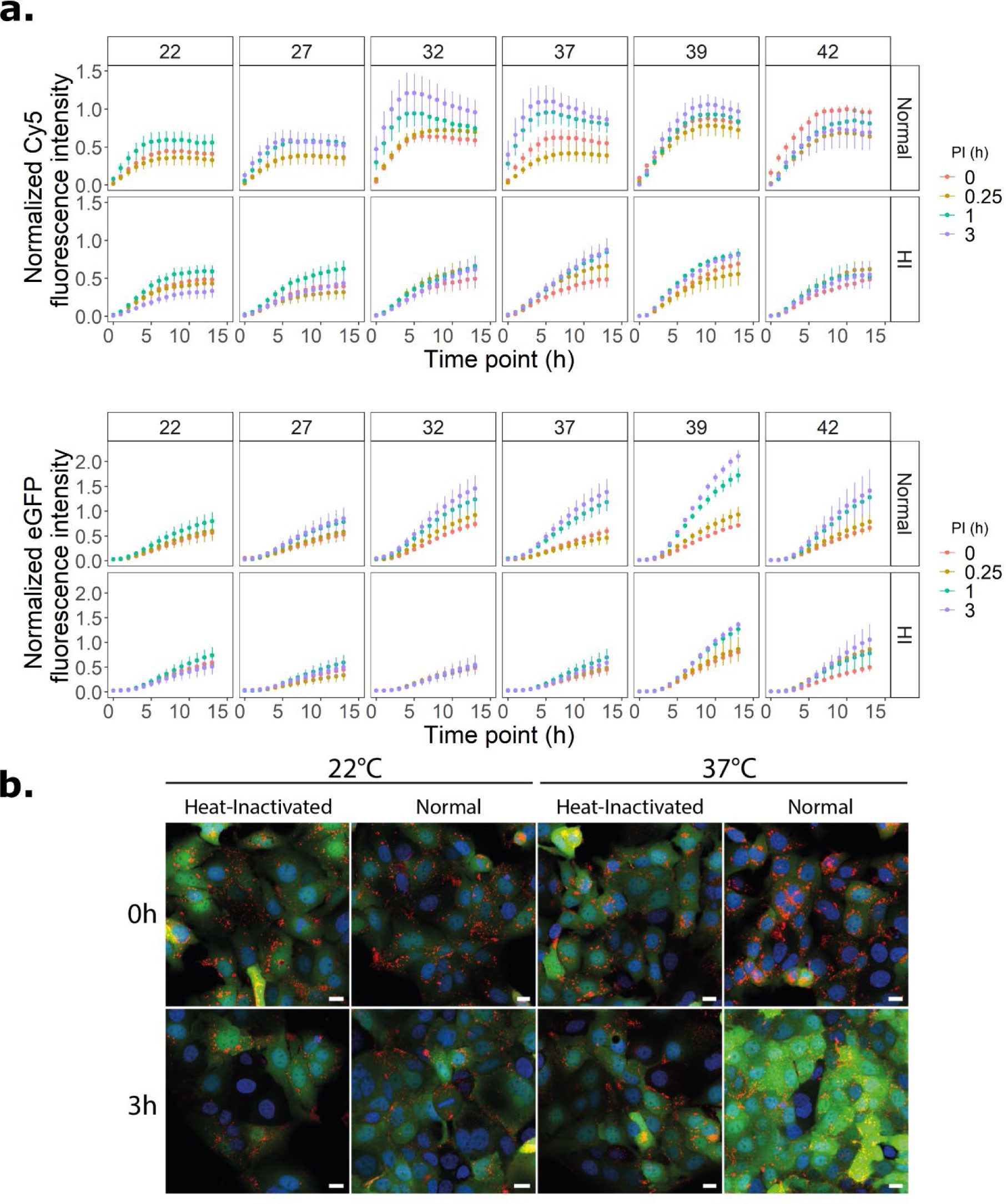
Effects of temperature and serum heat-inactivation on the cellular uptake and delivery efficiency of serum pre-incubated LNPs. **(a)** Live cell imaging-based screening of the effect of temperature (22 to 42℃) on the cell uptake (top, Cy5 signal) and protein expression (bottom, eGFP signal) following pre-incubation (PI) of LNPs in cell culture medium supplemented with normal or heat-inactivated (HI) FBS for 0-3 hours (see legends). Error bars are standard deviations of the sampling distribution from four technical replicates of two biological replicates, performed in two independent experiments. **(b)** Representative images from the screen showing LNP uptake (Cy5, red), protein expression (eGFP, green) and nuclei staining (blue). Scale bars: 10 µm. The experiment was conducted using MC3_DMPE_PEG LNPs at a concentration of 0.625 µg/mL (mRNA basis). Corresponding experiments with MC3_DSPE_PEG LNPs are shown in supporting **Figure S6**.

Focusing next on the effects of pre-incubation temperature and serum heat-inactivation on the uptake and protein production in Huh-7 cells exposed to the MC3_DMPE_PEG LNPs (**Figure 2**), we found that cell uptake (Cy5) was most effective when LNPs were pre-incubated at near physiological temperature (32, 37, and 39°C) in normal FBS. Pre-incubation in heat-inactivated FBS significantly extended the lag time behaviours reported in **Figure 1c-e**. This is consistent with previous literature showing that heat-inactivated serum reduces the uptake of both unmodified[49] and PEG-ylated[50] polystyrene nanocarriers, and suggests that both protein composition and protein conformation is important in the formation of an uptake-promoting protein corona. Notably, several variants of apolipoprotein E (ApoE), which has been reported as an important mediator of functional *in vivo* delivery of MC3-carrying LNPs[37] have melting temperatures in the heat-inactivation relevant range,[51] suggesting they may be partially denatured. At the lowest (22-27°C) and highest (42°C) temperatures there were no clear-cut effects of the pre-incubation time in normal FBS on cell uptake (**Figure 2a, top panels**), although at 42°C a longer pre-incubation time did result in higher eGFP production (**Figure 2a, lower panels**). This may be due to temperature-sensitive variations in both the adsorption propensity and binding affinity of corona proteins as reported by Mahmoudi *et al.* in a study of dextran coated FeOx nanoparticles.[52] Although the assayed temperature range is extreme in relation to the human body average (37℃), the observations of changes in LNP behaviour with temperature are important since body temperature, and hence the efficacy of an LNP therapy, can be expected to fluctuate, due to fever, physical activity, hormonal rates etc.[53–55] as well as inhomogeneities between cells within the body.[56, 57] Our data are thus in line with other findings on the effect of temperature and serum heat-inactivation on the uptake of solid nanoparticles, showing that the coronation of LNPs follow similar principles, but complement previous work by demonstrating the impact on protein-LNP interaction on the functional delivery (i.e. protein expression mediated by these LNPs).

### 2.3. Biophysical characterisation of serum-induced PEG-shedding and protein corona formation on LNPs

In order to identify the sequence of events that underlie the formation of an uptake-competent protein corona on the surface of LNPs, we used a combination of biophysical methods, including total internal reflection fluorescence (TIRF) microscopy, small angle neutron scattering (SANS) and NMR diffusometry to probe, individually or in combination, events of PEG-shedding and protein coronation *in vitro*.

First, we set up a TIRF-based assay to image individual LNPs, tethered to a functionalised surface via streptavidin-biotin coupling to a biotinylated PEG-lipid (0.006 mol%; corresponding to 10-20 biotin molecules per LNP) (**Figure 3a, supporting Table S1**). Upon addition of FBS-containing media, we observed, by TIRF time lapse imaging, detachment of individual LNPs from the surface (**Figure 3b**), indicative of PEG-shedding. The rate and extent of particle detachment depended on the concentration of FBS (**Figure 3c**). Notably, when the LNPs were incubated with 10% FBS, a pronounced acceleration in the rate of particle detachment was observed after ∼20 minutes. The kinetics of the process appears specific to the LNPs since corresponding serum-induced detachment of POPC liposomes doped with the same PEG-lipid occurred on a longer time scale (∼100 minutes, **Figure S6**). The results indicate that PEG-shedding is an early step in the series of events that underlie the formation of a serum-induced uptake-promoting protein corona on the surface of LNPs (compared to kinetics of uptake and protein expression in **Figure 1c, e**).

**Figure 3.**
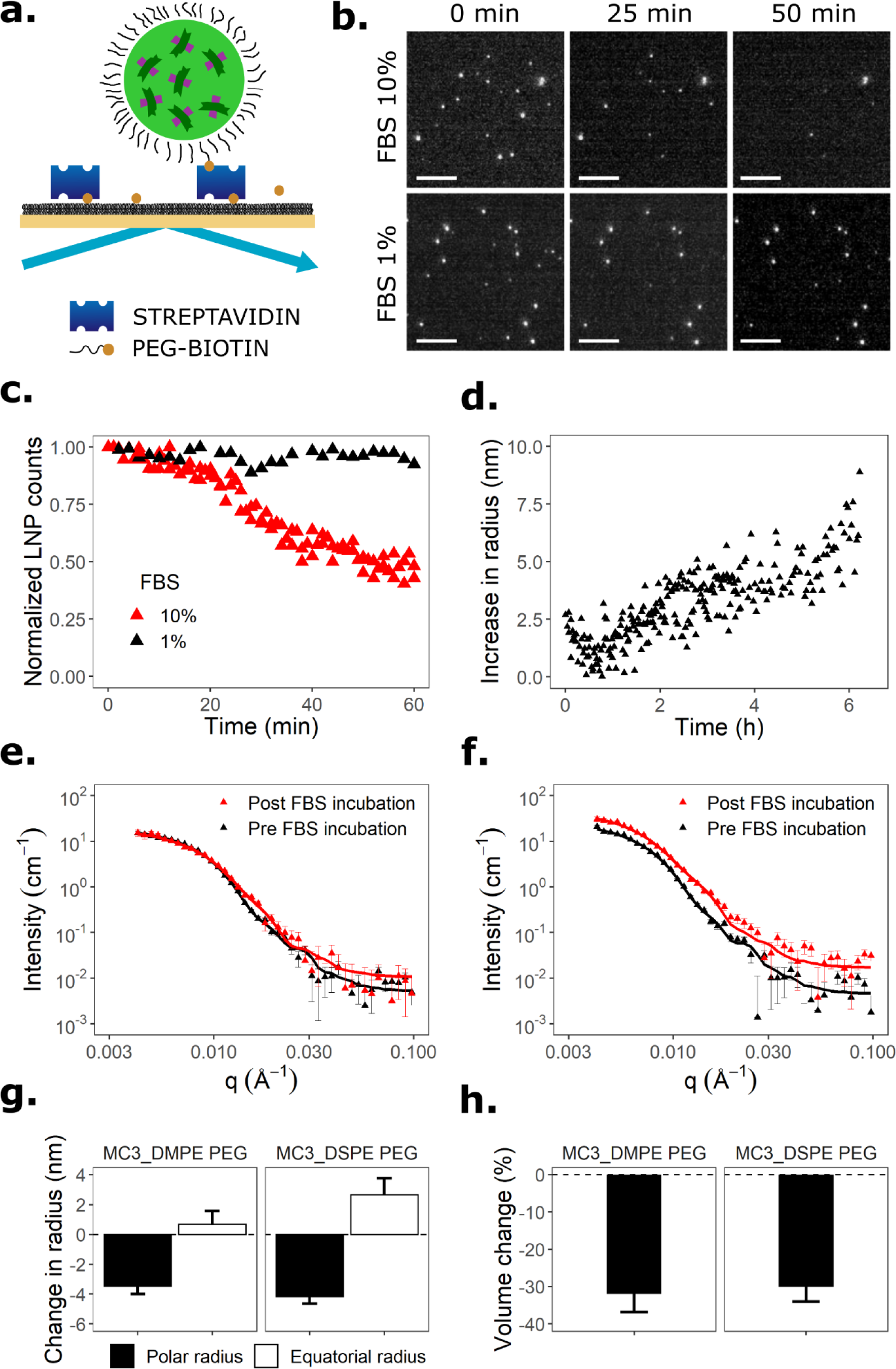
Biophysical characterization of serum protein induced PEG-shedding and coronation of LNPs. **(a)** Schematic illustration of the imaging-based approach to quantify detachment of single LNPs tethered to a supported lipid bilayer via a biotin-modified PEG-lipid (DSPE-PEG2000 biotin) binding to streptavidin. **(b)** Temporal changes in the number of surface-tethered LNPs quantified using the assay described in (a) following exposure to cell culture media containing different concentration of FBS (scale bars; 4 µm). **(c)** Representative images captured during the measurements in (b) showing the change in surface-tethered single LNPs over time. The experiments in (b) and (c) were performed at ambient temperature. **(d)** Time-dependent change in the hydrodynamic diameter of FBS-exposed LNPs determined by multimodal analysis of dynamic light scattering (DLS) autocorrelation function data. **(e-h)** Small angle neutron scattering (SANS) data showing the effect of FBS on (e) MC3_DMPE_PEG LNPs and (f) MC3_DSPE_PEG LNPs. The LNPs were diluted 10x in deuterated PBS and measurements were taken, at 37°C, pre- and post- incubation in PBS with 10% FBS for 3.7 hours in total at 37°C. (g) Change in the equatorial and polar radii of the LNPs obtained from fitting SANS data using an ellipsoid model. (h) LNP volume changes calculated from the equatorial and polar radii in (g).

To put the timing of PEG-shedding, as measured by LNP detachment above, in the context of protein corona formation kinetics, we used DLS to monitor broadly the effect of serum on LNP size (**Figure 3d and S7**). Since the analysed samples contained a complex mixture of large (**Figure S7a**; LNPs) as well as small-to-medium (**Figure S7b**; serum proteins in cell culture medium) components, we used multimodal analysis and followed overtime the LNP peak (∼ 100 nm) after mixing LNPs into cell culture medium (**Figure S8 and S7c**). The data suggest that the LNPs increase in size as function of time in serum. A rapid increase (delta radius = 2.5 nm) was observed during the first 5 minutes, supporting the early formation of an initial protein corona, followed by a continuous increase by > 5 nm in radius after 5 hours (**Figure 3d**). After 24 hours, the average diameter of the LNPs had increased by 30 nm, which may to some extent also result from particle aggregation (**Figure S7c**).

The effect of serum proteins on the structure of the LNPs was further investigated using small angle neutron scattering (SANS), comparing the standard MC3_DMPE_PEG (**Figure 3e**) to MC3_DSPE_PEG (**Figure 3f**) to explore potential differences due to PEG-shedding. Parameters obtained from fitted reduced SANS scattering patterns (**Figure 3e,f**) indicated that the LNPs undergo a structural change upon exposure to FBS. Complexation analysis using ScÅtterIV further verified that protein coronation had occurred (**Figure S8**). The SANS data were fitted using an ellipsoid model, demonstrating LNP sizes in good agreement with DLS data (**Table S2**). Following 3.7 hours of incubation in FBS (3 h incubation followed by 40 min measurement duration), we observed reduced polar radii and increased equatorial radii of both LNP types (**Figure 3g, Table S3**), consistent with volume reductions by 32.0 ± 4.9 % and 30.1 ± 3.9 % for MC3_DMPE_PEG and MC3_DSPE_PEG respectively (**Figure 3h**). We attribute this loss in volume to lipid dissociation, noting that the build-up of a protein corona will not be detectable as such, due to the lack of scattering contrast relative to the protein-rich background. Chen et al.[58] observed, using radiolabelled lipids, that LNPs with similar diameters (80 nm) and lipid compositions released approximately 60% of the DMG-PEG, and 15% of the DLin- MC3-DMA after 3 hours of incubation in mouse plasma. The volume reduction could also infer a structural rearrangement of PEG-chains. For DSPE-PEG2000 molecules incorporated into a DSPC lipid membrane, the difference in PEG layer thickness between the brush and mushroom conformations is approximately 3.5 nm,[59] which is well in line with the 27% volume reductions inferred from the measured SANS data.

To further explore the kinetics of PEG-shedding, we used nuclear magnetic resonance (NMR) diffusometry to measure, as function of time, the appearance of free DMPG_PEG in solution[41, 60] upon addition of FBS to MC3_DMPE_PEG LNPs (loaded with polyA RNA; **supporting Table S1**). Signal from the PEG lipids was identified at 3.58 ppm in the ^1^H NMR spectrum (**Figure S9a**), following the application of a water suppression method to decrease baseline from water (at 4.7 ppm) (**Figure S9b**). Titrations of LNPs with FBS verified no significant signal overlap between the PEG signal and the FBS (**Figure S9c**). Moreover, overlap with signals from residual ethanol (EtOH) from the LNP production were negligible at moderate gradient strength due to strong ethanol signal attenuation from fast diffusion. Figure 4a shows the normalized integrals of the PEG-signal from diffusion NMR plotted versus *k*=(*ygδ*)^2^(Δ-*δ*/3), recorded following the addition of 20% (v/v) FBS to LNPs in PBS/D2O; the corresponding data for free PEG-lipid in PBS/D2O is shown for comparison. The absence of a biexponential decay (first data points include EtOH contribution) for both free PEG and PEG in LNPs clearly indicates that the displacement exchange between the two environments is much faster compared to the total time (Δ) of the diffusion experiment. The low gradient strength of the probe forced the use of relatively large values of *δ* (5 ms) and Δ (200 ms) to acquire a sufficiently strong signal attenuation in order to calculate the self-diffusion coefficient. In the absence of FBS, the self-diffusion coefficient of PEG was low (weak slope) and corresponded to the expected diffusion of a spherical particle with a diameter of our LNPs (0.808e-11 ± 0.015e-11 m^2^/s), as verified by studying the NMR signals of cholesterol also belonging to the LNP. Upon addition of FBS, the slope increased (**Figure 4a**), indicating enhanced self-diffusion of released PEG-lipid as expected during PEG-shedding. The self-diffusion coefficient of released PEG-lipid, 3.56e-11 ± 0.04e-11 m^2^/s, is consistent with its assembly into micelles. The change in self-diffusion is rapid, and at the 27 minutes time point the data indicate a significant fraction of released PEG-lipid, which is consistent with the TIRF data (**Figure 3c**). Interestingly, after 30-40 minutes, the PEG-lipid diffusion coefficient (slope of the lines) starts to decrease to eventually equilibrate at a value corresponding to that of LNPs; this behaviour was not reported by Wilson et al. in their study of similar LNPs.[41] To obtain more detailed information on the PEG-shedding kinetics, we recorded real time diffusion-filtered one-dimensional spectra, with short time lapse, of LNPs in PBS/D2O, following the addition of 20% FBS at time zero. The resulting integral intensity of the PEG signal plotted versus time is shown in **Figure 4b**; corresponding values from Figure 4a are shown in red for comparison, showing the reproducibility of PEG-shedding. The data confirm that rapid and extensive PEG-shedding occurs during the first ∼30 minutes of FBS incubation. The consecutive increase of the PEG signal after 30 minutes, as discussed above, indicates that the shedded PEG-lipids, with time, either co-associate with FBS and serum proteins into larger particles or re-adsorb onto the surface of the LNPs. Notably, we could observe a time-dependent change in self-diffusion coefficient of PEG-lipids in FBS also in the absence of LNPs, suggesting that serum-induced self-association is possible.

**Figure 4.**
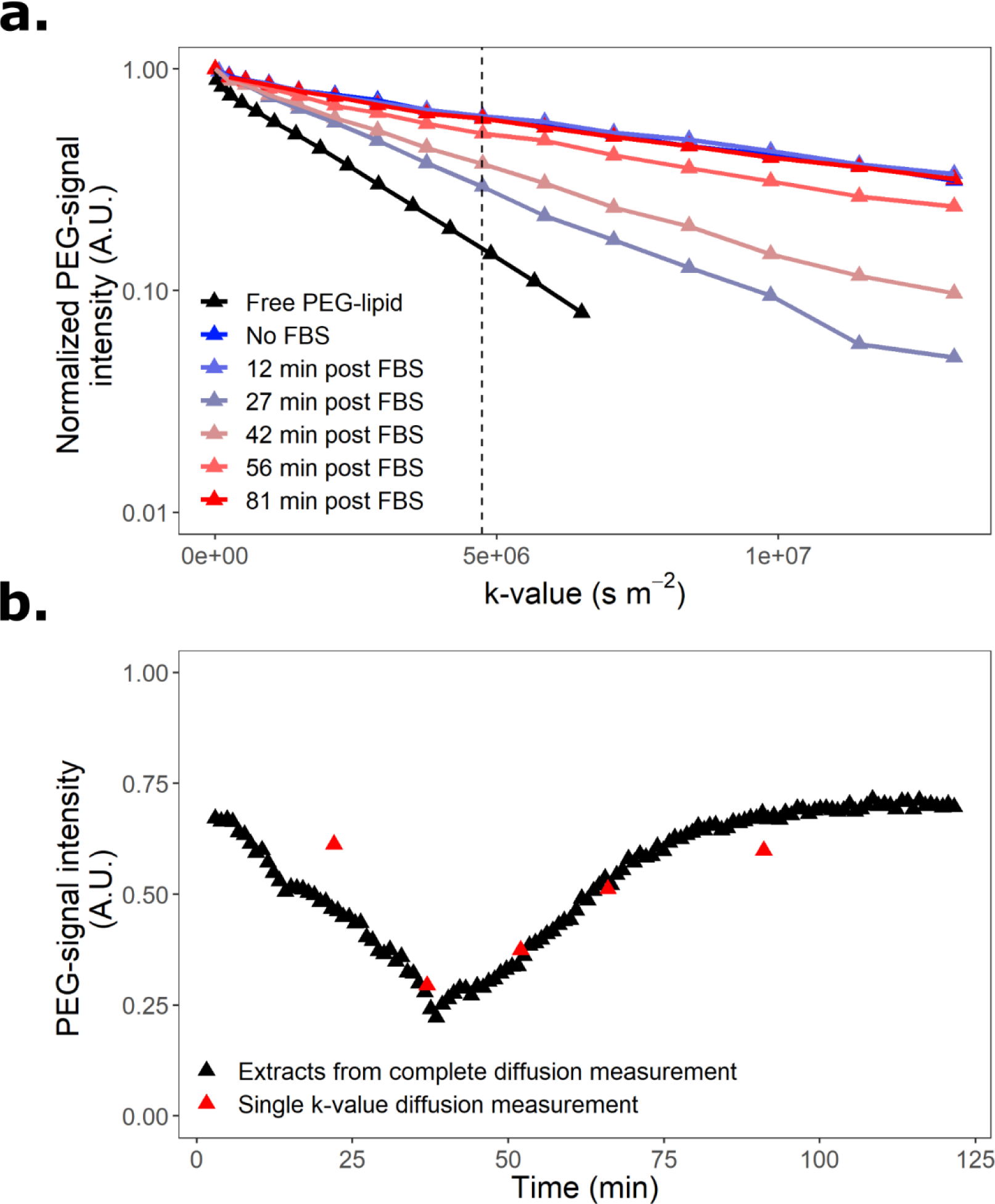
Kinetics of serum-induced PEG-shedding monitored by nuclear magnetic resonance (NMR) diffusometry. **(a)** Normalized integrals of the PEG-signal as function of k-value taken from a diffusion-series of LNPs incubated in PBS/D2O, with 20 % (v/v) FBS. Free PEG-lipid in PBS/D2O (black line) and untreated LNPs (no FBS, blue line) are included as reference. The initial intensity of the PEG-signal for each measurement is normalized to 1. **(b)** Intensity of the PEG-signal from a single *k*-value of 4.74·10^10^ s·m^-2^ diffusion-series of LNPs in PBS/D2O plotted against time after the addition of 20% FBS (v/v).

### 2.4. Characterisation of protein corona formation using quantitative proteomics

To explore the biological composition of the serum-induced protein corona forming on the LNPs, we performed an LC-MS/MS-based proteomics analysis, comparing the protein composition on MC3_DMPE_PEG LNPs harvested immediately upon dilution into cell culture media with 10% FBS (denoted 0 h) and after 1, 4, or 24 hours. Prior to analysis, unbound proteins were separated from the protein-LNP complexes using magnetic beads cross-linked with an anti-PEG antibody (IgM with KD=8.24 pM) able to recognize the backbone of the PEG molecule. After the pull-down, between 10 and 30 % of the LNPs were recovered. Protein digestion was thereafter performed on concentration matched LNP samples to ensure equal treatment efficiency. A multiple test (ANOVA, permutation based false discovery rate 5%) of the proteomics data led to the identification of 15 unique proteins for which significant changes in protein corona abundance were observed with time. The heatmap in **Figure 5a** displays the hierarchical clustering of the abundance of these proteins from the Z-score of Label-Free Quantification (LFQ) intensities. The LFQs are based on the intensities normalized on multiple levels to ensure that profiles of LFQ intensities accurately reflect the relative amounts of the selected protein across the samples.[61] Three distinct clusters were identified: cluster I groups proteins that rapidly interact with the surface of LNPs, such as Apolipoproteins C-III and A-II; cluster II groups proteins that preferentially enrich on the surface between 1-4 hours of pre-incubation, such as Apolipoproteins E and F; and cluster III groups proteins that enrich on the LNP surface after 24 hours, such as apolipoproteins B and M, and vitronectin. We also calculated the percentages of the intensity Based Absolute Quantification (iBAQ) values (**Figure S10**) for the 15 proteins, to rank their relative molar ratio within each sample.[62] This analysis showed that apolipoproteins A-II and E (ApoA-II and ApoE) are the two most abundant proteins associated with the MC3_DMPE_PEG LNPs, together accounting for between 69-82 mol% of all proteins present. Interestingly, we found that ApoE exhibited maximum abundance (of 40±3 %) after 4 hours (**Figure 5b**), conspicuously coinciding with the fastest LNP uptake (**Figure 1b**). This is important since ApoE, which plays a major role in the clearance and hepatocellular uptake of physiological lipoproteins, has been pointed out to have a crucial role for the *in vivo* delivery efficacy of MC3-containing LNPs in hepatocytes, and reported to mediate LNP uptake via the low-density lipoprotein (LDL) receptor.[45, 58, 63] All other proteins that were observed to adsorb preferentially to the LNPs at the 4 h time point had iBAQ percentages of 0.1% or lower, which indicates that their presence are less likely to be decisively important. Our data thus suggest that a time-dependent protein corona rearrangement to accommodate an enrichment of ApoE could underlie the lag time in uptake of LNPs reported in **Figure 1**.

**Figure 5.**
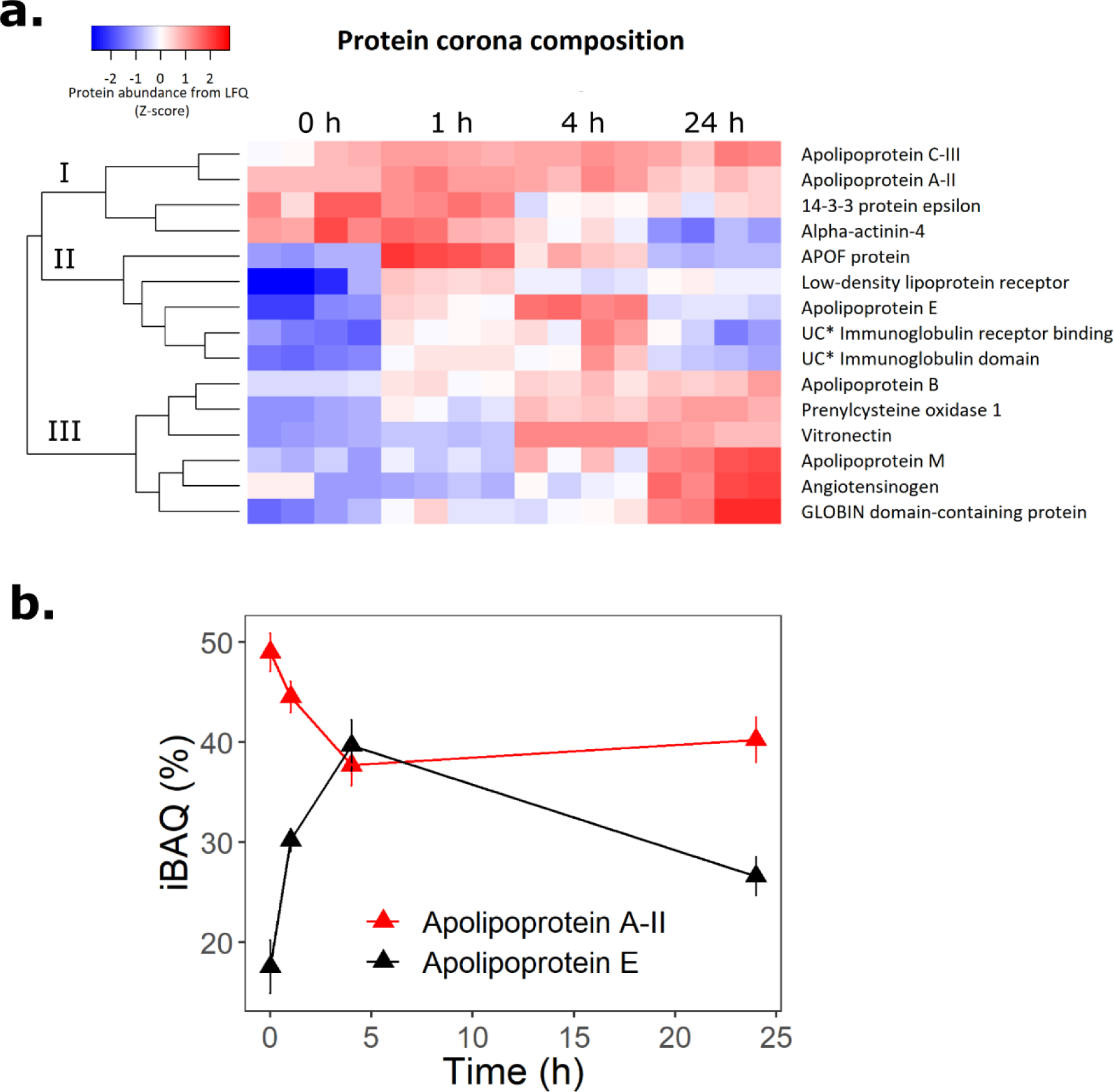
Characterisation of the serum-induced LNP protein corona by LC-MS/MS. **(a)** Heatmap with hierarchical clustering of the protein abundance (Z-score of Label-Free Quantification (LFQ) intensities) on MC3_DMPE_PEG LNPs that had pre-incubated in cell culture media with 10% FBS for different times (0 - 24 hours) and thereafter isolated from the medium using magnetic beads coated with an anti-PEG antibody. The heatmap displays data from two independent samples analyzed in duplicate (N=2, n=2). **(b)** Change in the percentages of the intensity Based Absolute Quantification (iBAQ) values of Apolipoprotein A-II and Apolipoprotein E, which were the two most abundant protein species on the surface of LNPs.

In contrast to ApoE, the abundance of ApoA-II, a common structural component of high-density lipoprotein particles, instead decreased with time (**Figure 5b**). This is interesting in relation to a previous observation of preferential adsorption to this lipoprotein to LNPs with poor siRNA delivery efficiency and enrichment of DSPE-PEG2000 lipids.[36] It is therefore possible that a PEG-shedding related partial release of ApoA-II from the LNP surface can help promote the uptake efficacy of our MC3_DMPE_PEG LNPs. In the class of proteins that enrich on the LNP surface over time and show highest abundance after 24 hours, and are thus likely constitute a component of the hard corona, we find for example vitronectin (**Figure S10**; maximum abundance 2.2±0.7 %) which has been shown to impede LNP-mediated delivery of siRNA to HepG2 cells.[37] This enrichment, in combination with the partial loss of ApoE after 24 hours in serum (**Figure 5b**), should be seen in light of our observations that LNPs that had been pre-incubated for this time period were taken up equally well as LNPs pre-incubated for 4 hours, albeit yielding only about half the eGFP protein expression (**Figure S5**). It is possible that the exact nature and composition of the protein corona is not only important for cell surface interactions (receptor binding) and uptake, but perhaps also the potency of the endosomal release of the mRNA and consequently its translation.

### 2.5. Serum pre-incubation of LNPs enables pulse-exposure and intracellular particle tracking

The principle finding of this study is that pre-incubation of LNPs in serum induces PEG- shedding and protein coronation, thereby enabling the LNPs to adopt uptake competent states. In addition to shedding new light on the nature, kinetics, and relative order of the different molecular events of this biologically crucial process, our study also paves the way for a new methodology to enable pulse-chase monitoring of LNP uptake in live cells. This makes possible to disentangle temporally convoluted processes, thereby enabling studies of how LNPs are sorted and trafficked by the endolysosomal machinery upon entering into the cells; information which is crucial to understand the mechanisms of endosomal escape and the mRNA release that leads to functional protein expression.

Building on the data presented above, we set up pulse-chase experiments with pre-incubation of the LNPs (PI; 0, 30, or 60 minutes), pulse-exposure of Huh-7 cells (PE; 5, 10, or 20 minutes), followed by a rapid washing step and a chase period (60 minutes) (**Figure 6a**). Flow cytometry quantification at the end of the chase period shows, as expected, that the cellular uptake (Cy5) and eGFP expression increase with both the duration of the short pulse-exposure and the, considerably longer, serum pre-incubation time (**Figure 6b**). Importantly, we show that after pre-incubation in serum for ≥ 30 minutes, a short 5 minutes exposure pulse is sufficient to obtain clearly detectable (by flow cytometry) intracellular LNP levels and eGFP signals after the 1- hour chase period. This was further demonstrated by time lapse confocal microscopy, where real-time visualization of the first LNP uptake (Cy5) was observed within seconds and eGFP signals could be readily observed to build up an hour later (**Figure 6c; Supporting Movie S2**).

**Figure 6.**
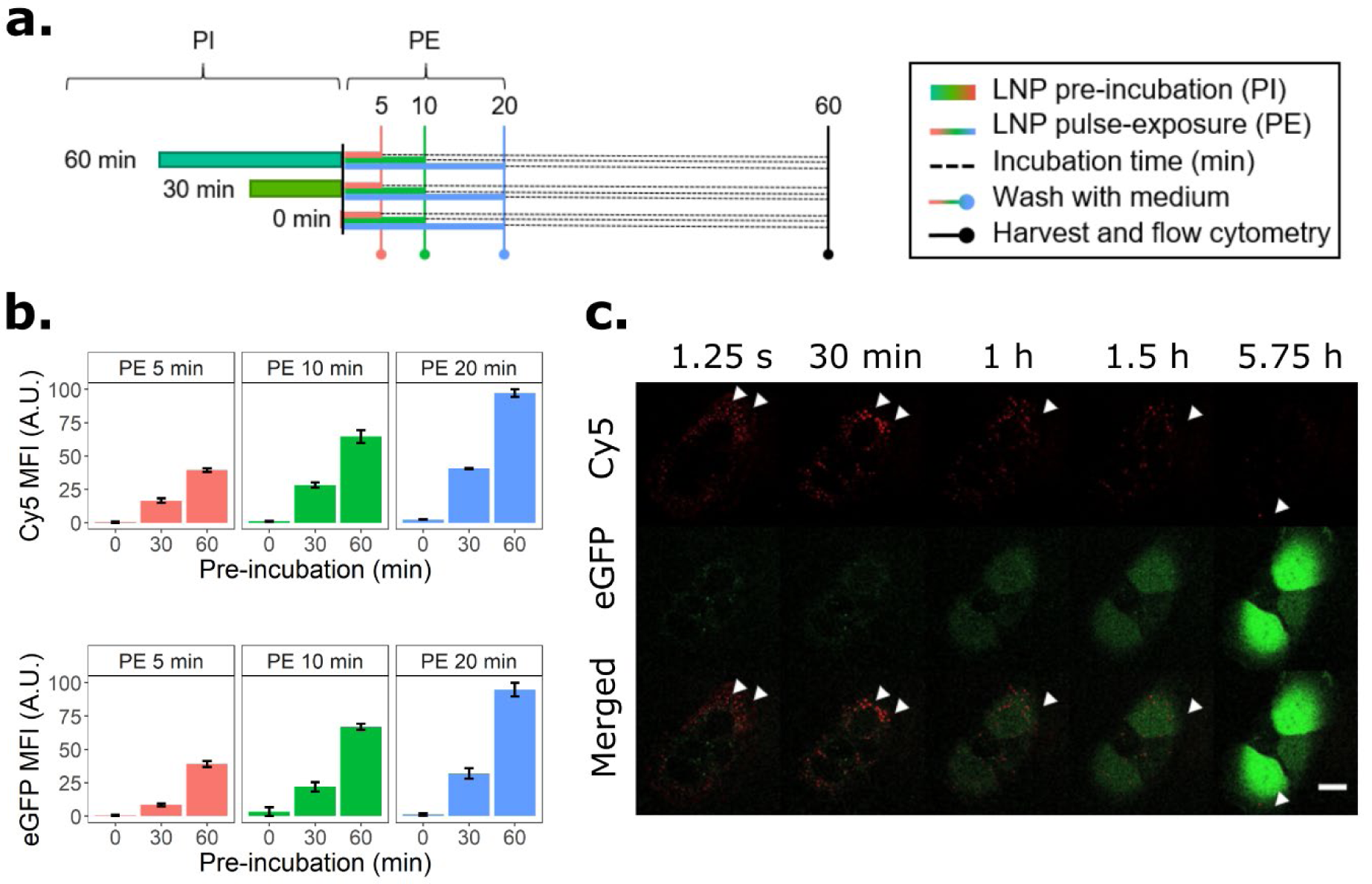
Serum pre-incubation of LNPs enables the monitoring of the cellular uptake and protein expression after short pulse-exposures (PE) of cells. **(a)** Schematic of the experimental setup used to quantify LNP uptake and protein expression using a combination of pre-incubation (PI) and short pulse exposure (PE) before monitoring the cells. **(b)** Quantification by flow cytometry of mRNA uptake (top, Cy5 signal) and protein expression (bottom, eGFP signal) as function of different pulse exposures (PE) and pre-incubation (PI) times. The LNPs were pre-incubated in cell culture medium with 10% FBS at 37°C. The LNP concentration was 1.25 µg/mL of mRNA. **(c)** Images from a live cell confocal microscopy time-lapse series recorded after pulse-exposure of Huh-7 cells for 5 minutes with LNPs (1.25 µg/mL of mRNA) that had been pre-incubated for 1 hour in cell culture medium with 10% FBS, (red: Cy5; green: eGFP; scale bar: 20 µm).

Using the pulse-chase strategy outlined in **Figure 6a** we then focused on probing the temporal co-localization of the MC3_DMPG_PEG LNPs with early endosomes (EE), late endosomes (LE) and lysosomes to monitor their intracellular trafficking. For this, we established Huh-7 cell models that were stably overexpressing mRFP-Rab5, mRFP-Rab7 or Lamp1-RFP to track each of these organelles, respectively. The endosome marker cell lines were cultured in separate chambers of a 4-well microscopy dish and simultaneously pulse-exposed for 5 minutes with LNPs that had been pre-incubated in serum-containing culture media for 2 hours. The cells were monitored by time-lapse imaging for 5 hours with a 30 second interval between image frames (**Movie S3-S5**). The image stacks were analysed as described in the Methods section to determine the fraction of Cy5-mRNA positive structures colocalized with RFP positives structures in each of the cell models and as a function of time (**Figure 7a**); the colocalization on which this image analysis was based is exemplified in **Figure 7b**, where white pixels in the colocalized pixel map corresponds to signal overlap. The data show that LNPs are rapidly trafficked through the endolysosomal system, with an initial preferential co-localisation with early endosomes occurring during the first hour post exposure. In this regard, our method allowed us to observe early colocalization between LNPs and Rab5-positive structures, on a time scale that agreed well with the onset of protein production (**Figure 6c**). The results highlight the importance of early endosomes as compartments that allow endosomal escape, and is well in line with published literature.[47, 48]

**Figure 7.**
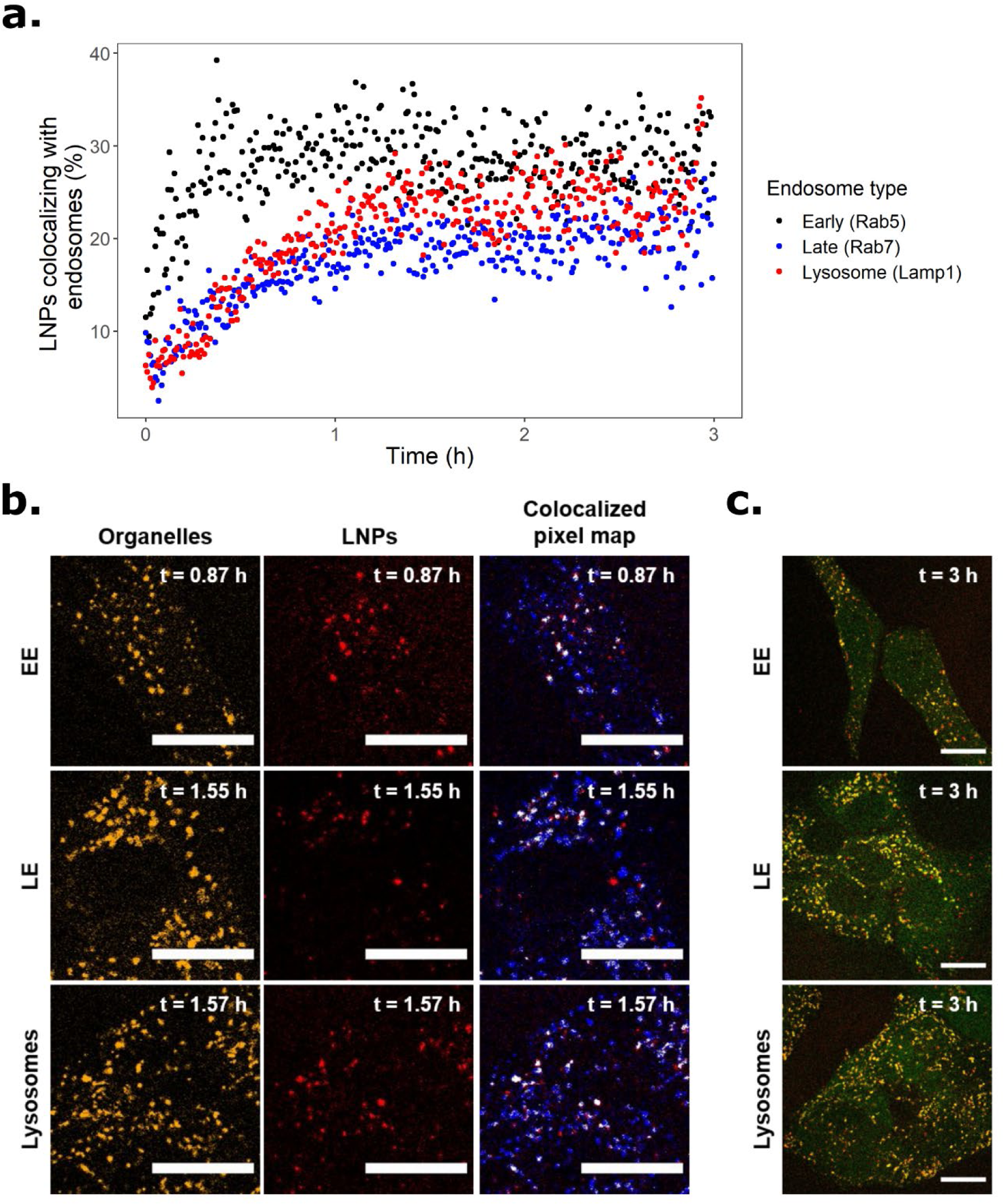
Time-lapse monitoring of the endolysosomal trafficking of LNPs following short pulse-exposure, enabled by serum pre-incubation of the LNPs. (a) Co-localisation of LNPs (Cy5 signal) with markers for early endosomes (mRFP-Rab5), late endosomes (mRFP-Rab7) and lysosomes (Lamp1-RFP) in Huh-7 cells following pulse-exposure for 5 minutes with MC3_DMPE_PEG LNPs (1.25 µg/mL of mRNA) that had been pre-incubated for 2 hours in cell culture medium with 10% FBS at 37°C, expressed as the percentage of LNP positive particles overlapping with the respective endosome marker as a function of time using the Pearson Correlation Coefficients (PCCs) between the two reporters. The data represent the means of two independent experiments (n=2) for each cell model. Each image frame typically contained 12-35 cells. The Huh-7 cells were genetically modified to stably overexpress the mRFP-tagged endosome marker proteins. **(b)** Representative confocal microscopy images (5.0x magnification) of the Huh-7 cells analysed in (a) showing endosomes and lysosomes as RFP positive particles (yellow) and LNPs as Cy5 positive particles (red). The colocalized pixel maps, obtained using the ImageJ Colocalization Threshold plugin, display the organelles in blue, LNPs in red and their overlap in white. (c) Confocal microscopy images (2.3x magnification) showing all signals detected, including eGFP protein expression, recorded 3 hours after pulse-exposure. The scale bars in all images are 20 µm.

## 3. Conclusion

Efficient cargo uptake and release are key features of a functional mRNA delivery system. We have studied the uptake and delivery of mRNA formulated into MC3 LNPs, focusing on the role of serum proteins in driving the temporal evolution of PEG-shedding reactions and protein coronation events that are necessary to prime these LNPs for fast and effective cell uptake. By studying the kinetics of LNP internalisation into Huh-7 hepatic cells under continuous exposure conditions, we show that their uptake is preceded by a significant lag time (∼2 hours in 10% FBS), during which PEG-shedding and protein coronation must take place. We also demonstrate that this lag-time can be effectively eliminated if the LNPs are pre-incubated in FBS. Even though serum interactions in the human body will differ kinetically and compositionally from the ones observed here, this study demonstrates a mechanistically important concept and the need to consider the timing needed for LNPs to reach their most uptake competent states. In this in vitro model scenario, the most efficacious LNP particle with respect to uptake formed first after 3-4 hours of pre-incubation in serum. This constitutes a significant delay between administration and delivery.

The finding that pre-incubation of LNPs enable them to adopt a protein corona that significantly enhance their uptake kinetics and eliminates the uptake lag time in cell experiments is of significant analytical relevance, which we demonstrate herein by showing that pulse-chase exposure with short (5 minutes) pulses is made possible only if the LNPs are pre-incubated in serum. This unlocks new opportunities to monitor the intracellular trafficking of LNPs, and their temporal co-localisation with different endosomal organelles. Our results point out early endosomes as important compartments during endosomal escape in line with previous suggestions[47, 48] and may, combined with new multiparametric approaches to quantify endosomal rupture events,[30] represent a significant step forward towards further optimisation of functional vehicles for mRNA uptake and release.

The combination of *in vitro* biophysical approaches and proteomics analysis used in this study to dissect the sequence of molecular events that prime MC3-LNPs for uptake has enabled us to propose a model for how serum interactions can drive the LNPs into their most uptake-competent state. We confirm, by comparison of shedding-competent MC3-DMPE-PEG2000 LNPs and shedding-incompetent counterparts (with DSPE-PEG2000, e.g. longer acyl chains)[40], that partial PEG-shedding is a key enabling event towards the formation of an uptake-competent LNP, as earlier proposed by Li et al.[39] We show, by both diffusion NMR and a purpose-designed TIRF based single nanoparticle detachment assay, that PEG-shedding kinetics are fast, taking place within the first hour of serum exposure. Proteomics analysis showed that this coincides in time with high abundance of the lipoprotein ApoA-II on the LNP surface. We propose that this lipoprotein, which is important for the formation of high density lipoprotein particles and thus has potential to complex lipids, as a potential mediator of the PEG-shedding reaction. Following PEG-shedding, the protein corona matures, both with respect to thickness and biological composition. We observed from SANS data recorded after >3 hours of serum exposure, that there is an overall decrease in lipid volume even in LNPs which, using other assays, do not demonstrate PEG shedding. A possible explanation for this is the re- configuration of non-shedded PEG-lipids from a brush to mushroom state during protein corona maturation that could contribute to the priming of LNP uptake by reducing the steric hindrance effect that is commonly associated with extended PEG-chains.[40] However, the observed volume reductions may also relate to the shedding of other lipid components as reported by Chen et al.[58] Notably, we observed that the lipoprotein ApoE is gradually recruited to the LNP surfaces during the first 4 hours of protein corona maturation, and that the maximum abundance of this important mediator of LNP uptake via the LDL-receptor[45, 63] coincides in time with the formation of maximally uptake-competent LNPs (e.g., with fastest and largest uptake).

In conclusion, our study has enabled a means to pin-point the order and kinetics of the molecular events that precede and prime LNP uptake. This has led to a mechanistic model that explains how rapid PEG-shedding, followed by relatively slow events of PEG-rearrangement and protein corona maturation, gradually drives the adsorption of large amounts of the uptake-promoting ApoE protein to the LNP surface, transitioning them into a state where their uptake can proceed without lag time, and with rapid kinetics (**Figure 8**). The knowledge generated in this model study shows that it is important to account for how LNPs kinetically interact with their biological environment and that this may even become a critical design parameter in the development of new RNA delivery vehicles adapted to respond optimally in different serum profiles and in response to serum variations which may occur in vivo. In conclusion, the comprehensive approach undertaken in this work may inspire new formulations and designs of nucleotide delivery systems, taking into account their biological interactions and coronation to obtain the improved functionality required for broad clinical use.

**Figure 8.**
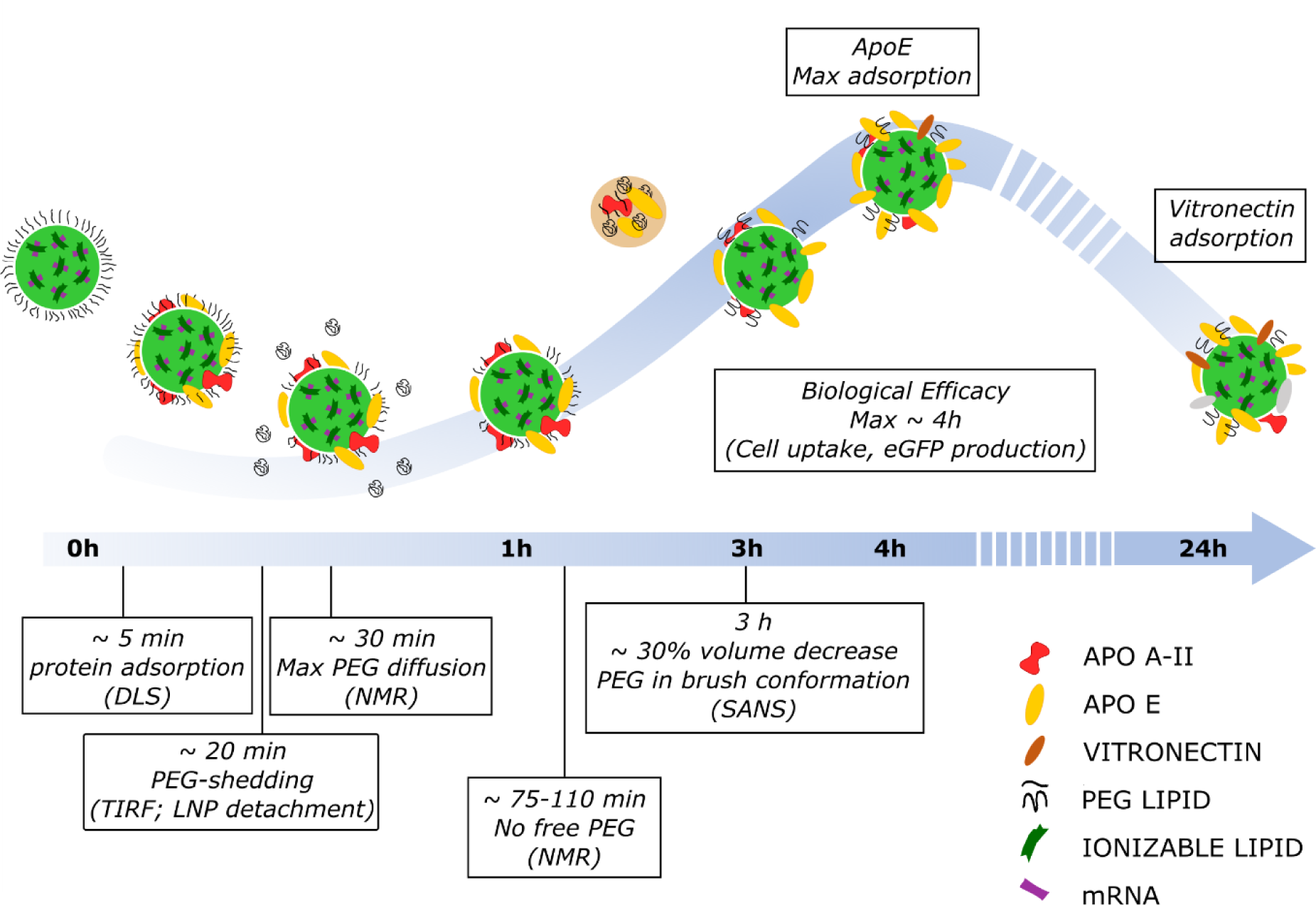
Proposed mechanistic model of PEG-shedding/rearrangement and protein corona maturation, driving biological efficacy of LNP.

## 4. Methods

### Material

The ionizable cationic lipid O-(Z,Z,Z,Z-heptatriaconta-6,9,26,29-tetraem-19-yl)-4- (N,N-dimethylamino)butanoate DLin-MC3-DMA (herein denoted MC3) was synthesized by AstraZeneca. 1,2-distearoyl-sn-glycero-3-phosphocholine (DSPC) and 1,2-distearoyl-sn-glycero-3-phosphoethanolamine-N-[biotinyl(polyethylene glycol)-2000] (DSPE-PEG2000- biotin) lipids were from Avanti Polar Lipids, 1,2-dimyristoyl-sn-glycero-3- phosphoethanolamine-N-[methoxy(polyethyleneglycol)-2000] (DMPE-PEG2000) and 1,2- distearoyl-sn-glycero-3-phosphoethanolamine-N-[Methylpolyoxyethyleneoxycarbonyl)-2000] (DSPE-PEG2000) were from NOF Corporation, and cholesterol from Sigma-Aldrich. CleanCap™ Cy5 eGFP mRNA (5-methoxyuridine) and CleanCap™ eGFP mRNA (5- methoxyuridine) (996 nucleotides) were from TriLink Biotechnologies. Polyadenylic acid (PolyA) and D2O for NMR, and streptavidin (used for the LNP detachment assay) were from Sigma-Aldrich. Citrate buffer (Teknova) and ethanol 99.5% was used in the LNP production; the mRNA was dissolved in HyClone HyPure Molecular Biology Grade RNase free water (GE Healthcare). DMEM cell culture media with high glucose, Glutamax supplement and pyruvate, Trypsin-0.25% EDTA, DPBS 1X calcium/magnesium free for cell culture, PBS (10X, pH 7.4) for dialysis, PBS tablets (-Ca^2+^, -Mg^2+^), the Quant-iT RiboGreen RNA assay kit, and Hoechst 33342 were purchased from Life Technologies. Fetal bovine serum (FBS) qualified Brazil (lot #10270106) from Life Technologies was used for all experiments.

### Formulation and characterisation of lipid nanoparticles (LNPs)

LNPs were prepared using a NanoAssemblr (Precision NanoSystems Inc.) for microfluidic mixing of lipids and RNA as described by Zhigaltsev *et al.*[64] The lipids were dissolved in ethanol at indicated molar ratios to a total lipid concentration of 12.5 mM (1.85 mg/mL). mRNA was diluted in RNase free citrate buffer (50 mM, pH 3.0). The two solutions were mixed in a 3:1 volume ratio at a rate of 12 mL/min to obtain LNPs with a mRNA:lipid weight ratio of 10:1 (MC3:nucleotide; 3:1 molar ratio).). The resulting LNPs were dialyzed overnight against sterile PBS pH 7.4 using 10 kDa cut-off Slide-A-Lyzer G2 dialysis cassettes (Thermo Scientific). For NMR, the LNPs were formulated with a PolyA cargo and concentrated after dialysis to ∼90 mg/mL of lipid by Amicon ultrafiltration with a 30 kDa MWCO (Thermo Scientific). Each batch of LNPs was characterised prior to use to determine particle size, polydispersity, particle concentration, and RNA encapsulation as described below.

### Dynamic light scattering (DLS)

The hydrodynamic (Z-avg) diameter of the LNPs was determined by DLS on a Zetasizer Nano ZS (Malvern Instruments Ltd) and at a lipid concentration of 45-50 μg/mL. The particle size distributions were calculated by cumulant analysis, setting the particle and dispersant refractive indeces to 1.45 and 1.33 respectively and the viscosity to 0.8872 cP at 25°C. DLS measurements of LNPs in presence of serum were performed at 37℃. Giving the complexity of the mixture (presence of serum proteins), the particle size was, in this case, calculated by multimodal analysis; the viscosity was set to 0.6864 cP. Three technical replicates per sample, each recorded by 11 consecutive recordings of 10 seconds each, were included in the analysis.

### mRNA encapsulation

The mRNA encapsulation was determined using Quant-iT™ RiboGreen™ RNA assay kit according to the manufacturer’s instruction. Briefly, LNP samples were diluted in Tris-EDTA (TE) buffer and assayed in either TE to determine non-encapsulated RNA or TE + 2% Triton X-100 to determine the total amount of RNA. 0.5% RiboGreen dye was added for read-out against a standard curve. The encapsulation was typically 97–99%.

### Nanoparticle tracking analysis (NTA)

NTA was used to determine particle concentration and as a second measure of LNP size. A NanoSight LM10 instrument equipped with a 488 nm laser and a Hamamatsu C11440-50B/A11893-02 camera (Malvern Instruments Ltd) and operating in scattering mode was used. The cameral level was set at 15 and detection threshold at 3, the movie collection time was 90 seconds and the flow rate set to 10 mL/min. All samples were measured in technical triplicate and at a concentration that yielded 50-100 particles in the field of acquisition. The recorded movies were analyzed with NTA 3.2 software (Malvern Instruments Ltd), setting the viscosity to that of water.

### Cell culture and exposure

Human hepatic Huh-7 cells (kind gift from Prof. Samir El-Andaloussi, Karolinska Institute) were cultured in cell culture media (CCM) containing DMEM high glucose, 2 mM L-Glutamine, 1 mM sodium pyruvate and 10% FBS. The cells were dissociated and passaged using calcium/magnesium free DPBS and Trypsin-0.25% EDTA. The medium was exchanged every three days during cultivation. The cells were tested and verified mycoplasma free. For experiments, the cells were seeded at a density of 0.18 million cells/mL one day prior exposure, unless else is noted. The seeded cells were exposed to LNPs that were either freshly diluted in cell culture media or pre-incubated with cell culture media with 10% FBS.

### Serum pre-incubation of LNPs

The LNPs were pre-incubated in CCM with 10% FBS at 37℃ and under 5% CO2. The LNP concentration was 0.625 µg/mL of mRNA (for continuous exposure) or 1.25 µg/mL of mRNA (for pulse-exposure). After pulse-exposures cells were washed 2X in FBS-containing CCM. For screening experiments exploring the effect of pre-incubation time, serum and temperature, LNPs were pre-incubated in CCM with 10% normal or heat-inactivated FBS at a concentration of 5 µg/mL of mRNA in a Veriti PCR thermal cycler (Applied Biosystems, California, US) and at temperatures between 22°C and 42°C.

### Flow cytometry

Cells were seeded in 48-well plates at a density of 250 µL/well. After LNP exposure, the cells were washed 2X with PBS, harvested with trypsin and treated with FBS-containing CCM for trypsin neutralisation. Samples were transferred to round bottom 96-well plates and analysed on a Guava® easyCyteTM 8HT (Millipore) flow cytometer. Cy5 and eGFP mean fluorescence intensities (MFIs) were recorded using excitation at 635 or 488 nm and emission detected at 661/19 nm or 525/30 nm, respectively. Data were processed using Flowing Software (version 2.5.1) (Turku Centre for Biotechnology) according to the following gating strategy; debris was excluded by gating on all events in the forward scatter (FSC) *vs* side scatter (SSC) plot for live cells; cell aggregates (doublets/clusters) were excluded by gating on live cells in a side scatter height (SSC-H) vs side scatter area (SSC-A) for single cells.

### Confocal microscopy

Cells were seeded in CELLview™ glass-bottom quartering cell culture dishes (Greiner Bio-One) at a density of 250 µL/compartment. The cells were imaged on an inverted Nikon C2+ confocal microscope equipped with a C2-DUVB GaAsP Detector Unit with variable emission bandpass, perfect focus system and an oil-immersion 60 × 1.4 Nikon APO objective (Nikon Instruments, Amsterdam, Netherlands) using a stage top incubator (OKOLab, Pozzuoli, Italy) for temperature and CO2 control. EGFP and Cy5-mRNA was excited sequentially by the 488 or 640 nm laser lines and fluorescence was detected between 496-566 nm and 652-700 nm for the green and red channel respectively. The pinhole aperture was set at 30 µm (1 Airy unit) unless else is noted. For LNP exposure in wild-type Huh-7 cells, confocal images were acquired for a total of 17 hours with 10 minutes interval in case of continuous exposure, or every minute for a total of 5.75 hours for pulse-exposed cells. To probe the intracellular trafficking of LNPs following pulse exposure, in-house established Huh-7 cells stably overexpressing mRFP-Rab5, mRFP-Rab7 and Lamp1-RFP were used, and time lapse movies recorded for up to 5 h with a 30 second interval between images, each field of view containing ∼12-35 cells. The fluorescence from the RFP reporter was excited by the 561 nm laser line and detected 569-636 nm. The Pinhole aperture was set at 90 µm (3 Airy units) to optimize particle tracking. All images were equally processed using Fiji (ImageJ 1.52p)[65] and colocalization analysis was performed using the EzColocalization plugin.[66] Briefly, the RFP channel was treated for background subtraction (rolling ball radius; 5) and uploaded as Ch.1, while Cy5 channel was not corrected and uploaded as Ch.2. All available colocalization analysis metrics were selected for the analysis and Pearson Correlation Coefficients (PCCs) between the two above reporters were displayed over time. The Colocalization Threshold plugin was used to generate colocalized pixel map displaying the organelles in blue, LNPs in red and their overlap in white.

### High content imaging

Cells were seeded in CellCarrier Ultra-384 well dishes (PerkinElmer) at a density of 5.5 x10^4^ cells/mL (50 µL/well). Prior to LNP exposure, cells were washed 2X and incubated in phenol red free DMEM with Glutamax containing 0.5 µg/mL Hoechst 33342 to stain nuclei. The pre-incubated LNPs (see above) were transferred to the cells using a Bravo liquid pipetting robot (Agilent Technologies, Santa Clara, US). All treatments were analysed in four technical replicates. Cells were then imaged on a CV7000 spinning disk confocal microscope (Yokogawa, Tokyo, Japan) in a humidified environmental chamber maintained at 37°C and supplemented with 5% CO2. The acquired images were analysed using Columbus software (version 2.9.1.532, PerkinElmer) to segment cellular structures and determine Cy5- mRNA uptake and eGFP expression. Data were normalized across experimental treatments to the 37°C control with 0 hours of pre-incubation in unmodified FBS.

### Total internal reflection microscopy (TIRF)

TIRF was used to monitor serum-induced detachment of LNPs tethered to the surface of CELLview™ glass-bottom quartering cell culture dishes (Greiner Bio-One) via a streptavidin-biotin linkage to DSPE-PEG2000 biotin lipids (0.006% incorporation). The LNPs were tethered by pre-incubation of the glass dish with streptavidin (1 ng/mL in PBS, 5 minutes) to achieve the low coverage density needed for single particle resolution. The surface was then washed 5X times in PBS and MC3_DSPE PEG-biotin LNPs (diluted in PBS to ∼ 6.5 e+9 particles/mL) were added. After 5 minutes, the surface was washed again 5X in PBS to remove unattached LNPs. The detachment of LNPs from the surface in presence of serum was then monitored for at least 90 minutes with TIRF videomicroscopy (1 minute frame rate, 200 ms exposure time) on a Nikon Eclipse Ti-E inverted microscope with a CFI Apo TIRF 100x (NA: 1.49) oil immersion objective and perfect focus system (PFS) (Nikon Instruments, Amsterdam, Netherlands), a Lumencor Spectra X LED light source (Lumencor Inc., Beaverton, US) and an Andor iXon EM+ DU-897 EMCCD camera (Andor Technologies, Belfast, UK). The LNPs were detected by Cy5 mRNA emission using an F46- 006 ET filter set (Chroma Technology GmbH, Olching, Germany). The recorded data was analysed using a custom-written script in MATLAB R2017B (MathWorks, 2017, Natick, MA, USA).

### Small angle neutron scattering (SANS)

SANS was used to study structural changes to the LNPs following pre-incubation with 10% FBS. LNP samples were diluted in deuterated PBS to avoid incoherent scattering from H2O. Scattering measurements were collected at the ZOOM beam line of the ISIS neutron and muon source at the Rutherford Appleton Laboratory, Didcot, UK, between 16 and 20 µA proton current. 1mm Hellma quartz cuvettes thermostated at 37°C were used. The beamline was configured with L1 = L2 = 4 m collimation and sample-detector distances to give a scattering vector Q = (4π/λ)sin(θ/2) range of 0.004 to 0.8 Å^-1^, where θ is the scattering angle and neutrons of wavelengths λ of 1.75 to 16.5 Å were used simultaneously by time of flight. For serum pre-incubation, LNPs were mixed with 10% FBS diluted in deuterated PBS and incubated in Eppendorf tubes at 37°C for 3 hours prior to 40 min measurement duration. Data reduction was performed using Mantid [67] and scattering simulations fitted using SasView version 4.2.2 (http://www.sasview.org/). For FBS-incubated samples, the FBS background was subtracted prior to data fitting. An ellipsoid model was used to fit the LNP data as detailed in the Table S1. The scattering length density (SLD) of the solvent and LNPs was fixed in the analysis to respectively 5.7034 x10^-6^ Å^-2^ (solvent), 1.21 x10^-7^ Å^-2^ (MC3_DMPE_PEG), and 1.35 x10^-7^ Å^-2^ (MC3_DSPE_PEG). The polar and equatorial radii from the ellipsoid fitting were used to calculate the FBS-induced change in LNP volume. To verify that complexation between LNPs and serum proteins had occurred, complexation tests were performed using ScÅtterIV where the monomer units were defined as the untreated LNP and the FBS background, and the complex as the LNP scattering post-incubation (unsubtracted).

### NMR spectroscopy

PEG-shedding investigations were conducted using pulsed-field gradient NMR spectroscopy on a Varian Inova 500 MHz spectrometer operating at 11.7 T with a 5 mm HFX probe with a maximum field gradient strength of 60 G/cm. Single pulse experiments included an 8 µs ^1^H-pulse, 2 s acquisition time, and 5 s recycle delay. For the diffusion measurements, the DgcsteSL (DOSY Gradient Compensated Stimulated Echo with Spin Lock) pulse-sequence combined with a dpfgse (double pulsed field gradient spin echo) solvent suppression sequence, was used, as shown in Figure S9a. The latter sequential part, based on excitation sculpting, was optimized to effectively suppress the large water signal of the samples. For the PEG-shedding investigation LNPs, PBS, and D2O were added directly and mixed in the NMR tube. All measurements have been run at room temperature with 200 ul D2O in total solvent volume 600 µl, the other constituents depending on the sample measured. FBS was added immediately prior to placing the NMR-tube in the magnet. Two different time-lapsed series of diffusion-based spectra were acquired to study the PEG-shedding. Firstly, a complete diffusion measurement with 16 gradients steps and full signal attenuation was acquired to obtain the self-diffusion coefficients for the different species. Secondly, a one-dimensional measurement was acquired repeatedly, generating a series of diffusion-filtered spectra a one specific gradients strength to capture the kinetics of the diffusivity change in presence of FBS. For both diffusion measurements, 8 µs ^1^H-pulse, 2 s acquisition time, 5 s recycle delay, 8 scans per point, *δ* of 5 ms, and Δ of 200 ms was used. The calibration of the water suppression was performed using a regular single pulse experiment with dpfgse addition. Relevant parameters for the water suppression included a 2.3 ms soft ^1^H-pulse and 1 ms gradient pulse of 16.87 G/cm. For all experiments a gradient-90-gradient-delay steady-state pre-pulse sequence was used to kill of all residual magnetization from previous scans.

For the complete diffusion measurement, gradients in 16 steps ranging from 0.18 to 60 G/cm were used, corresponding to *k*-values of 1.24·10^6^ to 1.31·10^11^ s·m^-2^, where *k*=(*ygδ*)^2^(Δ-*δ*/3). In order to follow the PEG-shedding over time, the number of scans for each spectrum was carefully chosen to have a suitable compromise between noise level and time lapse between consecutive spectra. In order to obtain sufficient signal-to-noise ratio, 8 scans were acquired for each spectrum giving a time between each consecutive diffusion measurement of approximately 17 minutes.

For the single k-value diffusion-filtered spectrum a gradient strength of 36.56 G/cm, corresponding to a *k*-value of 4.74·10^10^ s·m^-2^, was used, as chosen from the complete series of diffusion measurements. The time-series of 1D spectra captures the entire PEG-shedding event. With the same number of scans chosen as in the full diffusion series, the time between consecutive spectra could be decreased to below one minute.

### Proteomics Analysis of LNP corona: particle pull-down and protein digestion

The biological composition of the serum-induced LNP corona was investigated using LC-MS/MS-based proteomics analysis. LNPs (1 µg/mL of mRNA, ∼6 e+10 particles/mL) were incubated in CCM with 10% FBS for 0, 1, 4 or 24 hours. Dynabeads™ M-270 Epoxy (Thermo Fisher Scientific) cross-linked with anti-PEG [PEG-2-128] antibody (Abcam, Cambridge, UK) were used to isolate the protein-LNPs complexes. Quantification of the LNP pull-down was performed by Cy5 fluorescence readout. Protein digestion was performed on concentration matched samples to ensure equal amounts of recovered LNPs. Protein denaturation and reduction were done in a 30 minutes one-step reaction using urea and (tris(2-carboxyethyl)phosphine), followed by a 30 minutes alkylation step using 2-chloroacetamide. Protein digestion was thereafter done overnight in trypsin and ceased by the addition of formic acid. The LC-MS/MS based analysis was performed using a Q-Exactive HF mass spectrometer (ThermoFisher) coupled to an Evosep One (Evosep) automatic sample loader with disposable C18 trap columns for peptide desalting and purification. Purified peptides were then separated on an 8 cm HPLC column (Waters) with gradient off-set focusing to achieve a 3%-40% acetonitrile within a 44-minute loop at a 0.5 µL/min flow rate. The raw data was analyzed by MaxQuant software (version 1.6.6.0) and peptide lists were searched against the Bovine Uniprot FASTA database (version June 2019) with cysteine carbamidomethylation as a fixed modification and N-terminal acetylation and methionine oxidations as variable modifications. The false discovery rate was set to 0.01 for both proteins and peptides with a minimum length of 7 amino acids and was determined by searching a reverse database. Enzyme specificity was set as C-terminal to arginine and lysine as expected using trypsin as protease, and a maximum of two missed cleavages were allowed in the database search. Peptide identification was performed with an allowed initial precursor mass deviation up to 6 ppm and an allowed fragment mass deviation of 20 ppm. Matching between runs was performed with complete serum, depleted serum and lipoprotein fraction. Proteins matching to the reversed database were filtered out. LFQ was performed with a minimum ratio count of 1. To obtain a complete description of the pattern of mean differences among the conditions, a pairwise ANOVA comparison was performed on biological and technical replicates. Changes were found significant when at least 1 condition was different from the others. Bioinformatics analyses were first performed with the Perseus software (version 1.6.2.3). Protein relative abundances was computed using peptide label-free quantification values. For data visualization and clustering analysis, a false discovery rate of 5% in multiple sample testing was used.

## Supporting Information

Supporting Information is available from the Wiley Online Library or from the author.

## Acknowledgements

This work has been performed within the Swedish Foundation for Strategic research (SSF)- funded Industrial Research Centre “FoRmulaEx” (IRC15-0065) through funding to E.K.E.. A.G. acknowledges additional support from “Helge Ax:son Johnsons stiftelse”, “Wilhelm och Martina Lundgrens Vetenskapsfond” and “Kungliga Vetenskaps- och Vitterhets-Samhället”. H.M.G.B. holds an Individual Marie Skłodowska-Curie Fellowship “SmartCubes” (703666). M. O. acknowledges support from the Jane and Aatos Erkko Foundation. M.N.H. acknowledges support of the Swiss National Science Foundation (P300PA_171540). M.M.S. acknowledges support from the grant from the UK Regenerative Medicine Platform “Acellular / Smart Materials – 3D Architecture” (MR/R015651/1) and the Rosetrees Trust. The Area of Advance Materials Science at Chalmers University of Technology is acknowledged for supporting the NMR study. We thank Prof. S. El-Andaloussi for providing the Huh-7 cell line, Dr R. Bordes, Chalmers, for fruitful discussions regarding NMR and M. Thomas and R. Tonkin for their assistance during the SANS beamtime. Experiments at the ISIS Neutron and Muon Source were supported by allocation to H.M.G.B. (principal investigator), M.N.H., M.O., and M.M.S. from the Science and Technology Facilities Council (RB 1920358). This work benefited from the use of the Mantid (2013) application. Mantid (2013): Manipulation and Analysis Toolkit for Instrument Data.; Mantid Project. http://dx.doi.org/10.5286/SOFTWARE/MANTID. This work benefited from the use of the SasView application, originally developed under NSF award DMR-0520547. SasView contains code developed with funding from the European Union’s Horizon 2020 research and innovation programme under the SINE2020 project, grant agreement No 654000.

## Supporting Information

### Lipid nanoparticle formulation

LNPs with an eGFP-encoding Cy5-labelled mRNA cargo were formulated using an automated microfluidic platform to ensure reproducibility between the batches used in the study (Figure S1). supporting Table S1 summarizes the different LNP formulations used in this work, including the corresponding lipid compositions and cargos, as well as the type of experiment conducted. As a standard formulation, we used mRNA-containing LNPs with a lipid compositions of DLin-MC3-DMA:DSPC:Chol:DMPE-PEG2000 in the mol% ratio 50:10:38.5:1.5 (referred as MC3_DMPE PEG in the manuscript) for biological experiments and protein corona study. The eGFP-encoding Cy5-labelled mRNA was mixed with corresponding eGFP-encoding non-labelled mRNA (ratio 1:4) to ensure optimal combination of particle detection (fluorescence) and translation efficiency. In addition, variants were formulated, such that the MC3_DMPE PEG-biotin with an incorporation of 0.006 mol% of a DSPE-PEG biotin linker, was used for the LNP detachment assay using TIRF. For NMR, the mRNA cargo of the MC3_DMPE PEG formulation was substituted to polyA RNA. The particle diameter was determined using dynamic light scattering (DLS) and nanoparticle tracking analysis (NTA). The hydrodynamic (z-avg) diameters extracted from DLS were found to be 84-88 nm and highly consistent among the batches with mRNA encapsulation, the polydispersity indexes (PDI) < 0.09 reflect monodisperse populations of LNPs. LNPs encapsulating polyA had a slightly larger particle diameter (101 nm) and were more polydisperse (PDI = 0.333). The size measurements by NTA, which relies on the Brownian motion instead of light scattering, were consistent with DLS data and moreover provided particle concentrations (supporting Table S1). The mRNA or polyA encapsulation efficiency was determined to 97-99% using the RiboGreen assay, corresponding to an mRNA concentration of 124-139 µg/mL and a polyA concentration of ∼123 µg/mL.

**Figure S1.**
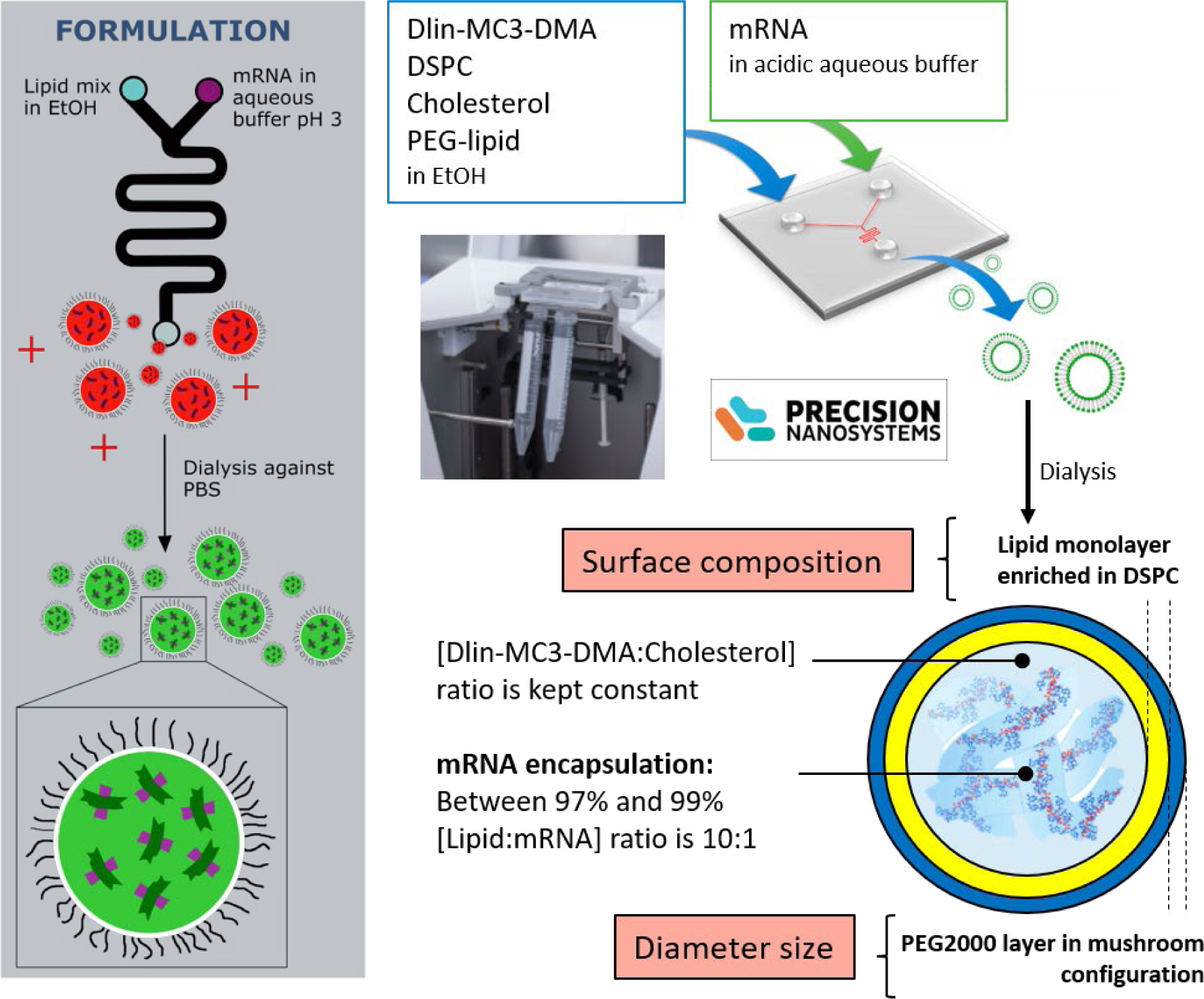
Schematic illustration of the LNP formulation using a microfluidic system to mix the lipid components (in ethanolic solution) with aqueous mRNA at pH 3.0. The resulting LNPs were dialysed against PBS to restore physiological pH.

**Figure S2.**
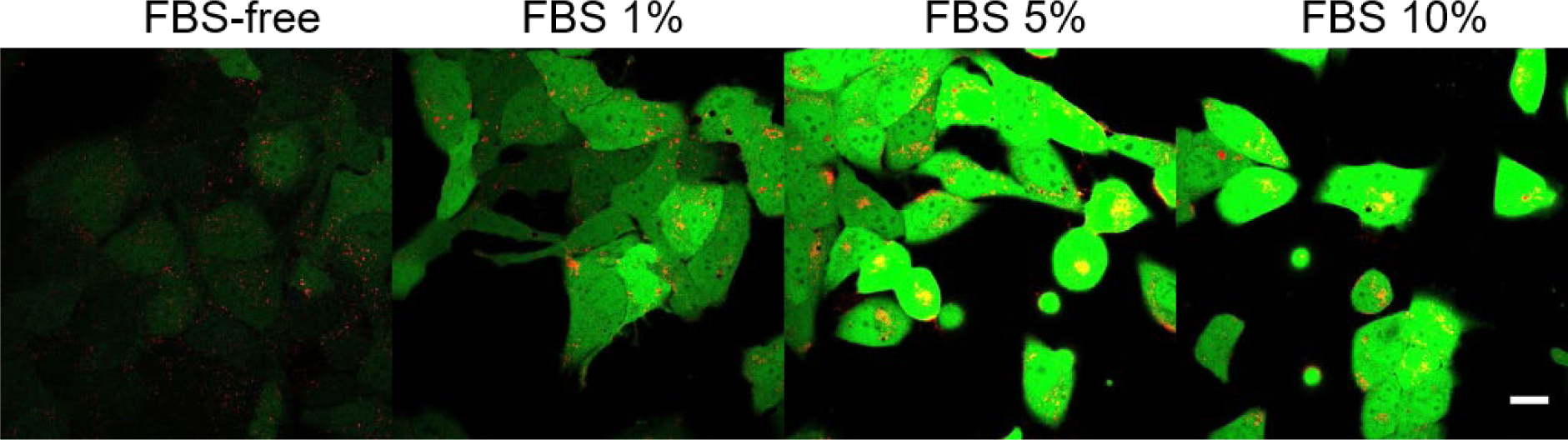
Serum-dependent uptake of LNPs after 17 hours continuous exposure. Confocal images of live Huh-7 cells exposed to MC3_DMPE_PEG LNPs (0.625 µg/mL of mRNA) for 17 hours in cell culture media with indicated concentrations of fetal bovine serum (FBS) showing overlaid cellular uptake (Cy5; red) and protein expression (eGFP protein; green. Scale bar: 20 µm).

**Figure S3.**
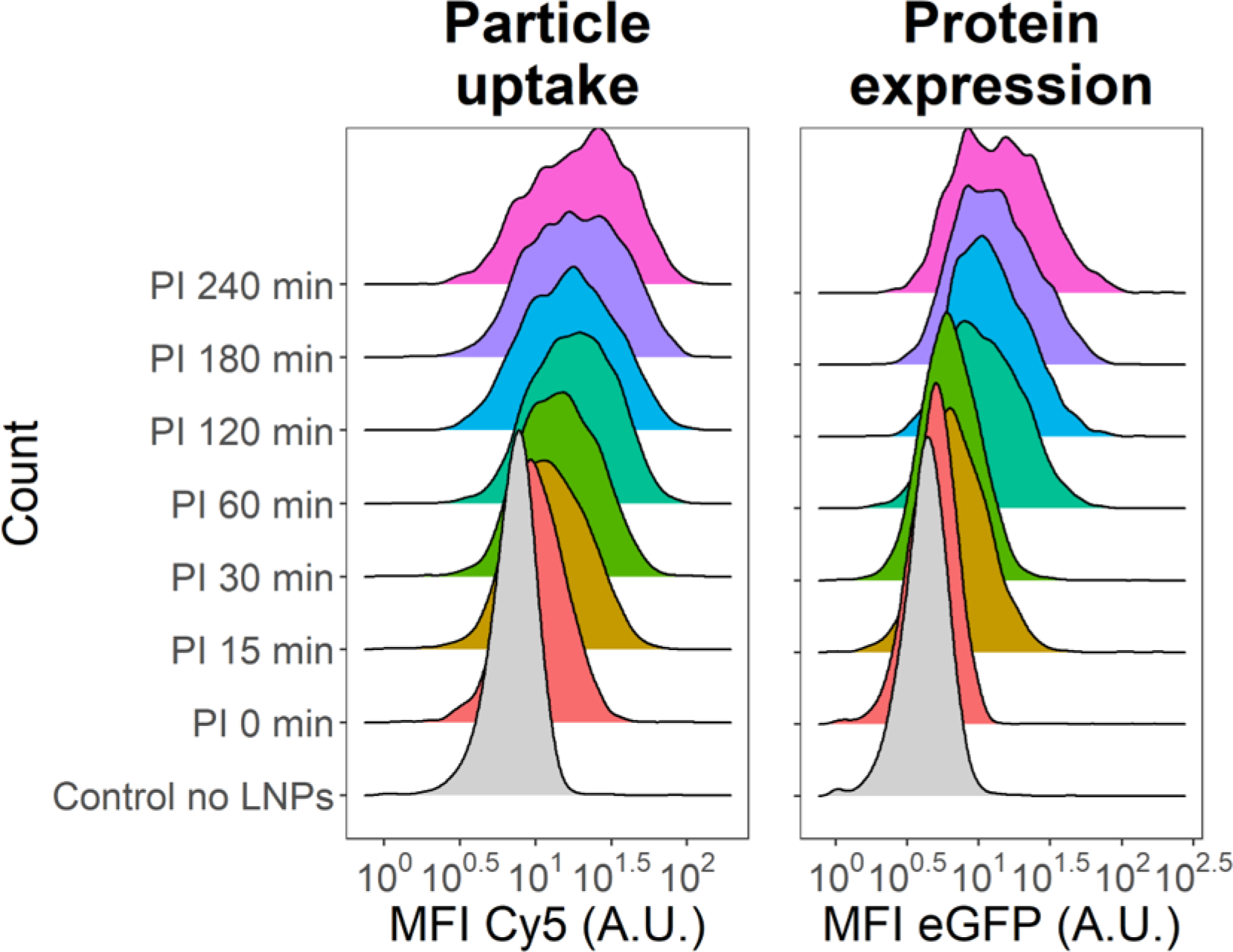
Cell uptake and protein expression 2 hours after exposure of Huh-7 cells with FBS pre-incubated LNPs. Representative histograms of the Cy5 and eGFP cellular mean fluorescence intensity (MFI) distributions across different pre-incubation (PI) times.

**Figure S4.**
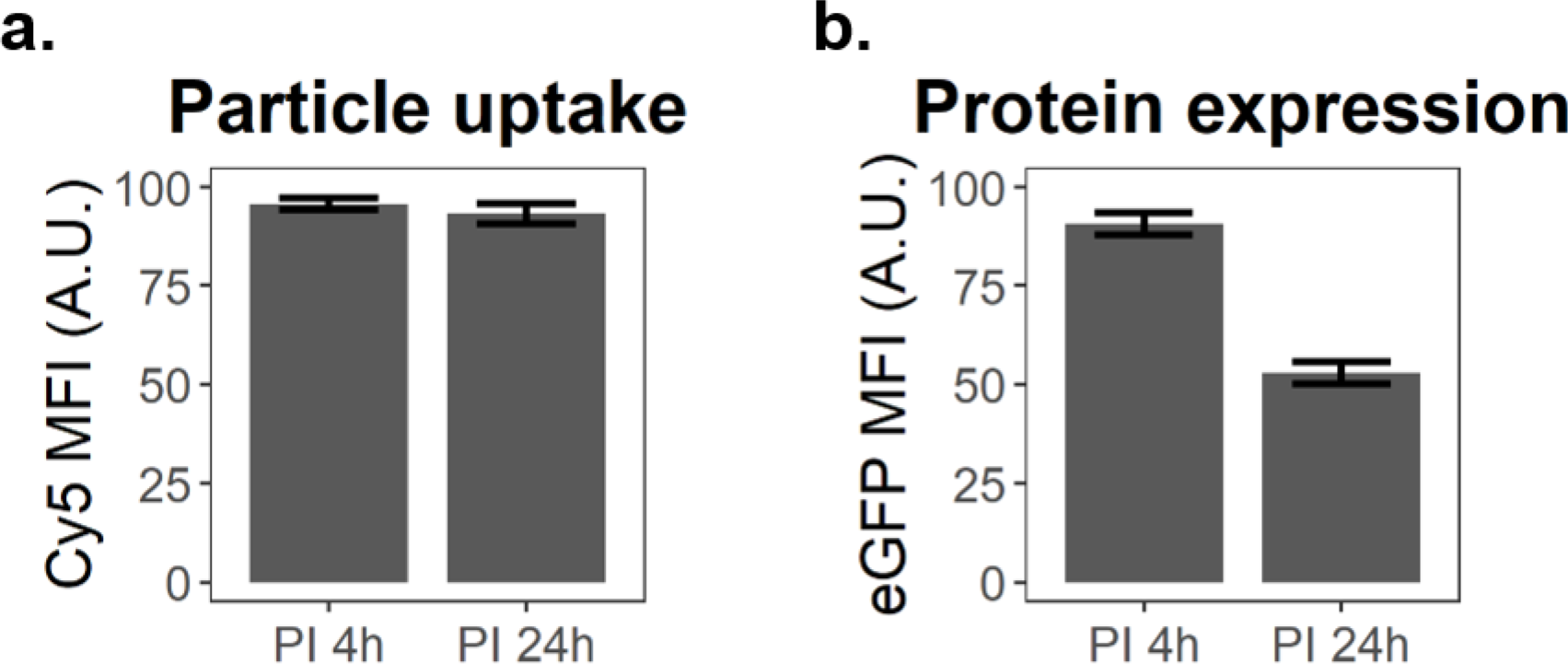
Optimum in the pre-incubation time for protein expression. Flow cytometry results displaying the (a) LNPs uptake and (b) eGFP expression after 2 h exposure with LNPs pre-incubated (PI) 4h or 24 h. Error bars are standard deviations of the sampling distribution from triplicate performed in two independent experiments.

**Figure S5.**
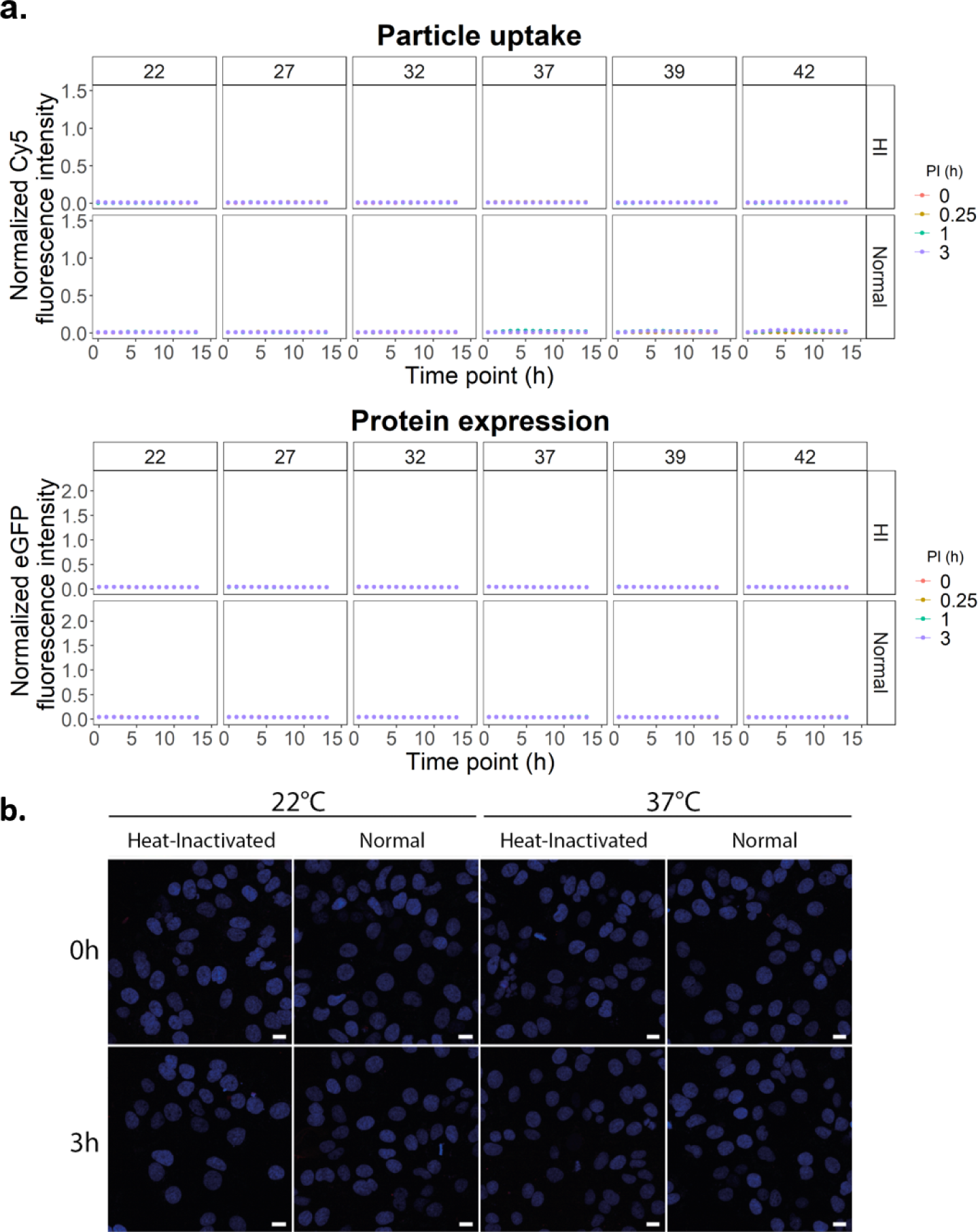
Effects of the temperature and serum heat-inactivation on delivery efficiency of pre-incubated LNPs using high-content imaging. Huh-7 cells exposed to pre-incubated MC3_DSPE_PEG LNPs in culture medium supplemented with Normal or heat-inactivated (HI) FBS for 3, 1 or 0.25 hours at temperatures from 22 to 42℃. (a) Live cell imaging of intracellular Cy5 and eGFP were detected overtime across the different conditions. (b) Representative samples of the analyzed images. Scale bars: 10 µm. Error bars are standard deviations of the sampling distribution from four technical replicates of two biological replicates.

**Figure S6.**
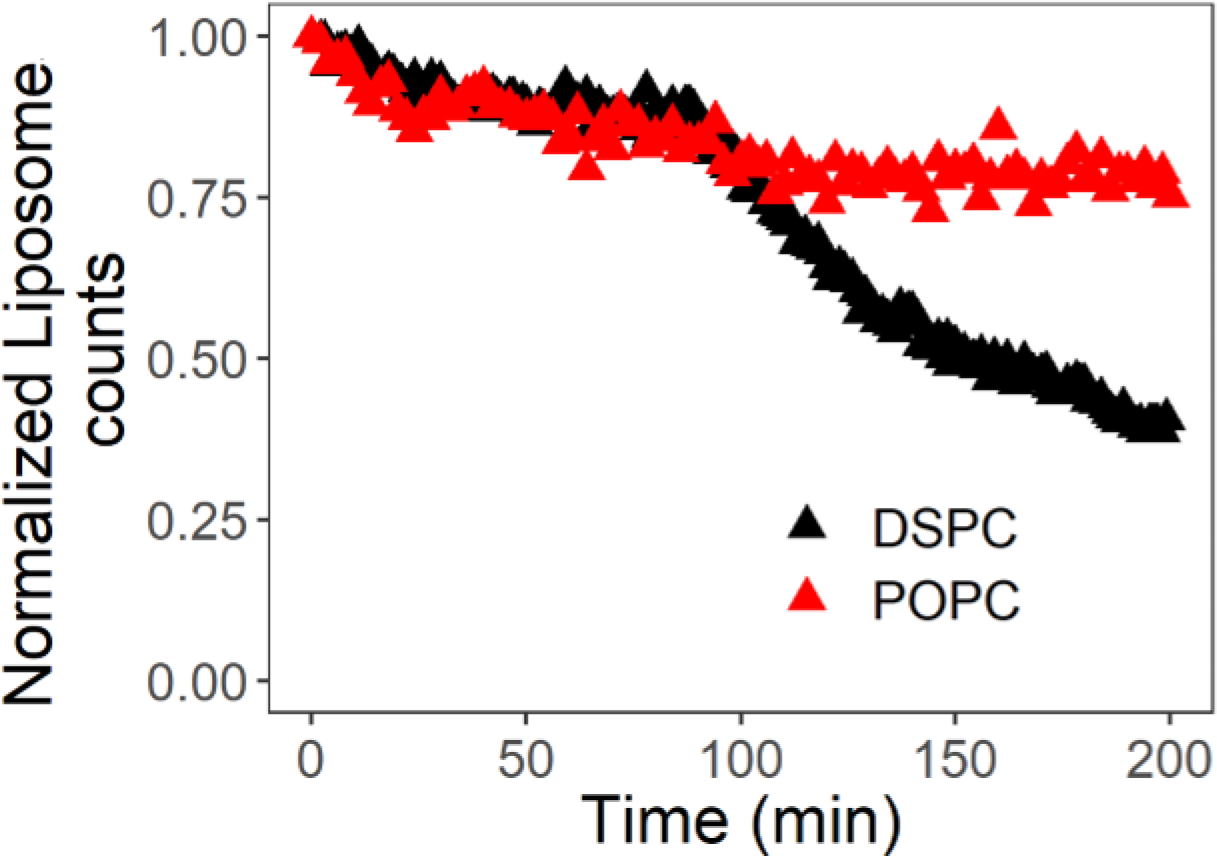
Kinetics of detachment of surface-tethered liposome in response to FBS containing media. Fluorescently labeled, zwitterionic-PEGylated (5 mol %) fluid-phase POPC and gel-phase DSPC liposomes were prepared using the extrusion method, with a size of approximately 100 nm, which is similar to that of LNPs. A small fraction of biotinylated PEG lipid (1-2 biotin per liposome) was also included for tethering the liposome to SA-coated glass surface. After incubation with cell culture media containing 10% FBS, a lag-burst kinetics was observed for detachment of POPC liposomes with a lag time of around 100 min (the lag time of LNPs was around 30 min). However, DSPC liposomes remained attached to the surface during the time course of the experiment (4 hours). The results show that the primary determinant of the rate of particle detachment is the lipid composition.

**Figure S7.**
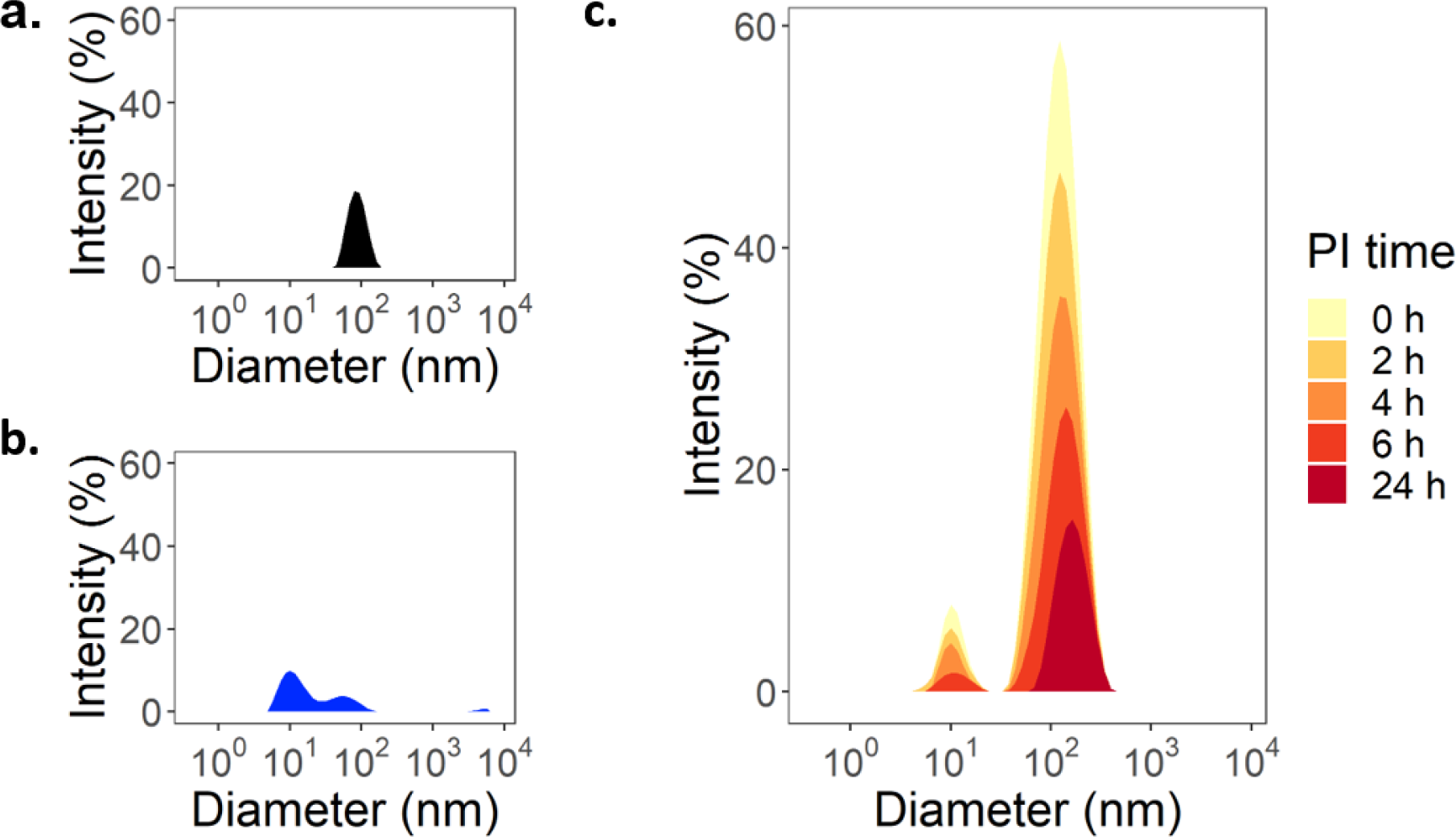
Representative spectra of LNPs diameter size measurement. DLS representative recorded spectra for (a) LNPs in PBS, (b) cell culture medium alone and (c) LNPs in cell culture medium across pre-incubation (PI) time.

**Figure S8.**
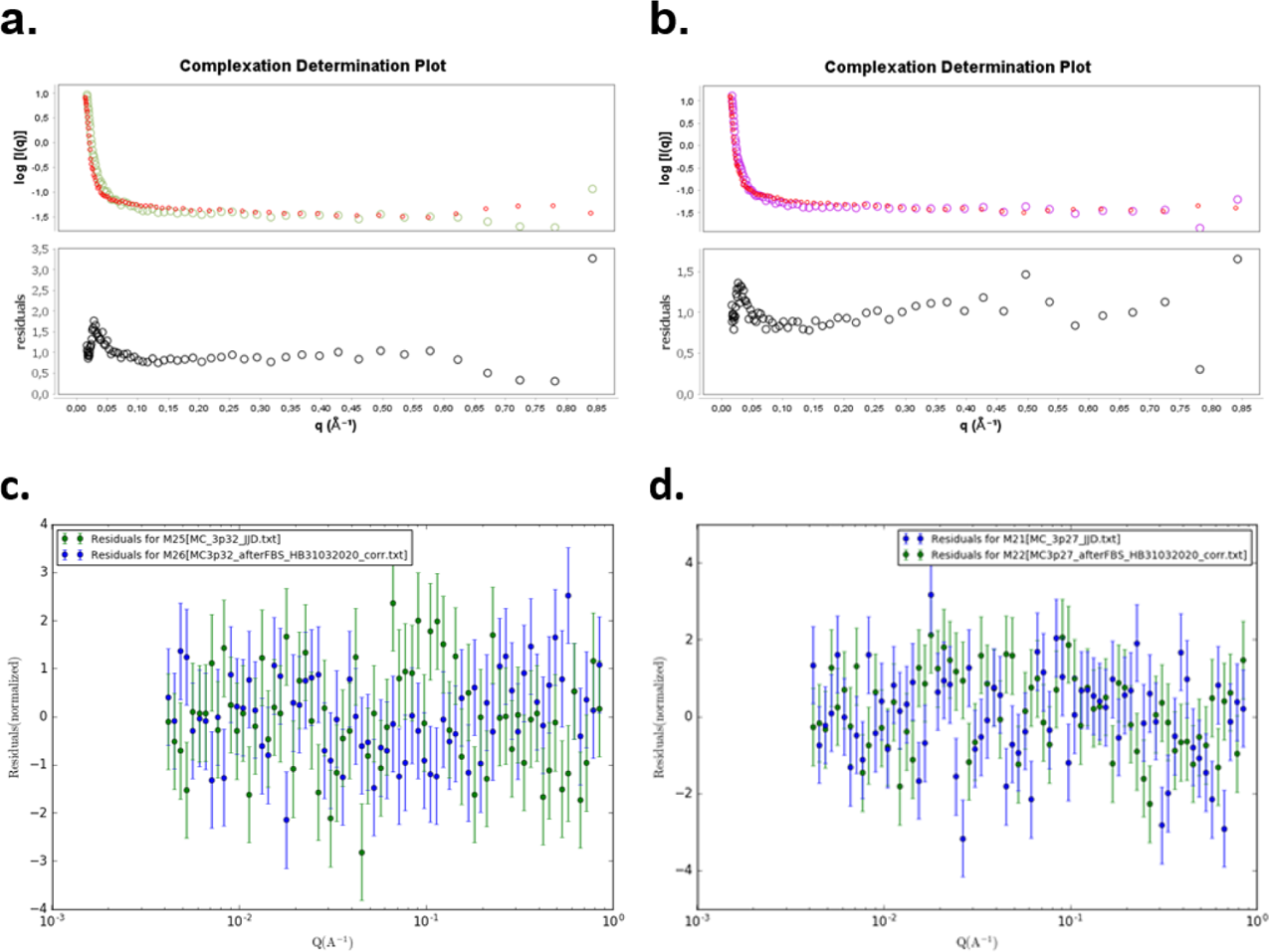
Residual plots from ScÅtterIV complexation analysis and SASView 4.2 fitting. The residuals in (a) MC3_DMPE PEG and (b) MC3_DSPE PEG, exported by ScÅtterIV using the complexation tool, indicate that the SANS scattering curve post FBS incubation is not a linear sum of the scattering curves obtained from the LNPs pre-incubation and the FBS background i.e. complexation has occurred. The residuals in (c) MC3_DMPE PEG and (d) MC3_DSPE PEG were exported from SASView 4.2 after fitting using the ellipsoid model.

**Figure S9.**
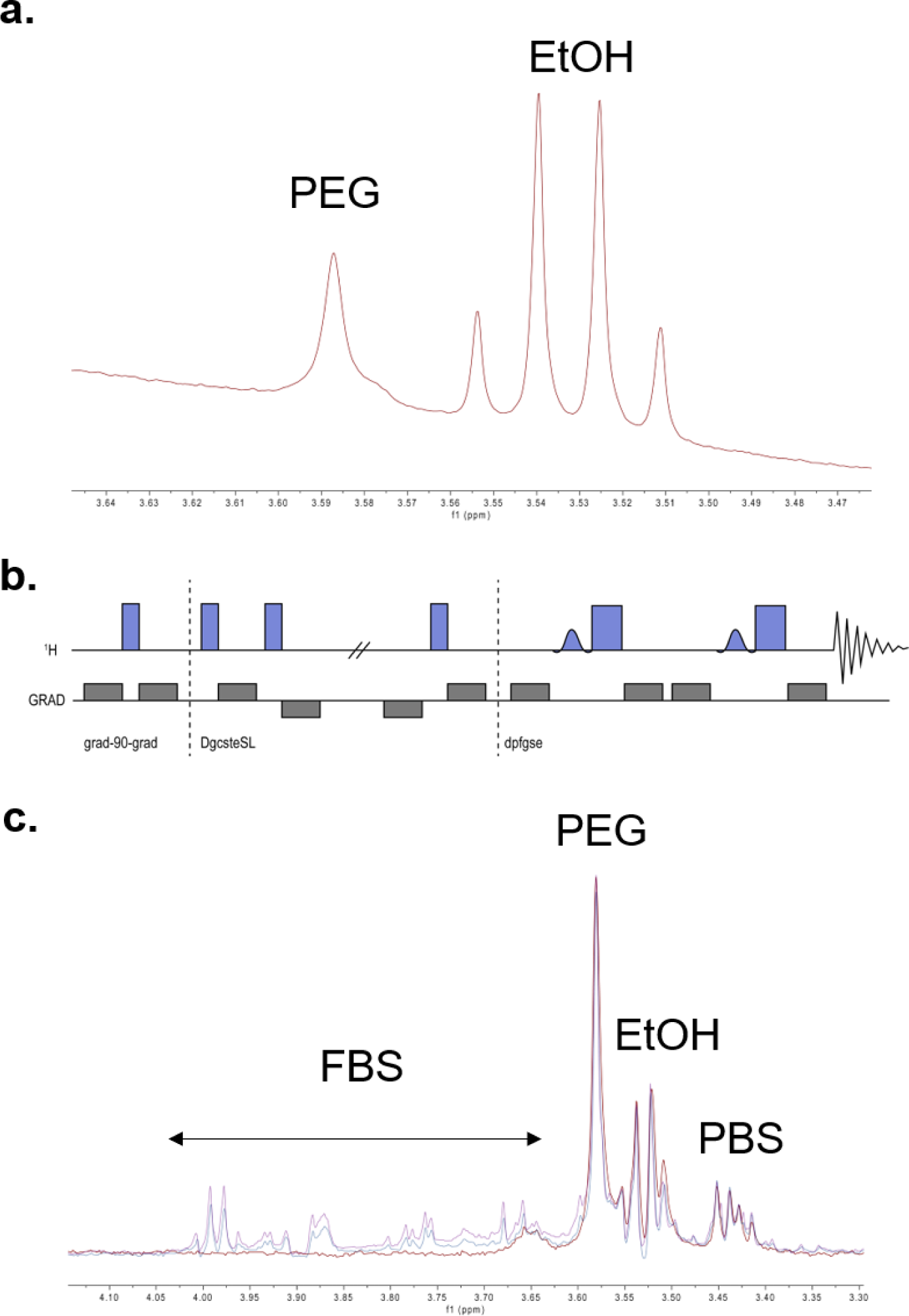
Nuclear magnetic resonance (NMR). (a) Part of ^1^H-NMR spectrum of sample with LNPs in D2O showing the signal from the PEG-lipid as well as residual EtOH in the sample. The impact from the extremely large water signal at 4.7 ppm can clearly be seen as a tilted baseline. (b) Schematic representation of the DOSY Gradient Compensated Stimulated Echo with Spin Lock (DgcsteSL) pulse-sequence used to suppress large water signals. (c) First point in the diffusion series of samples with LNPs in PBS/D2O, with 0% (red), 10% (blue), and 20% (purple) FBS (counted on the PBS). The different constituents of the samples are marked in the spectra.

**Figure S10.**
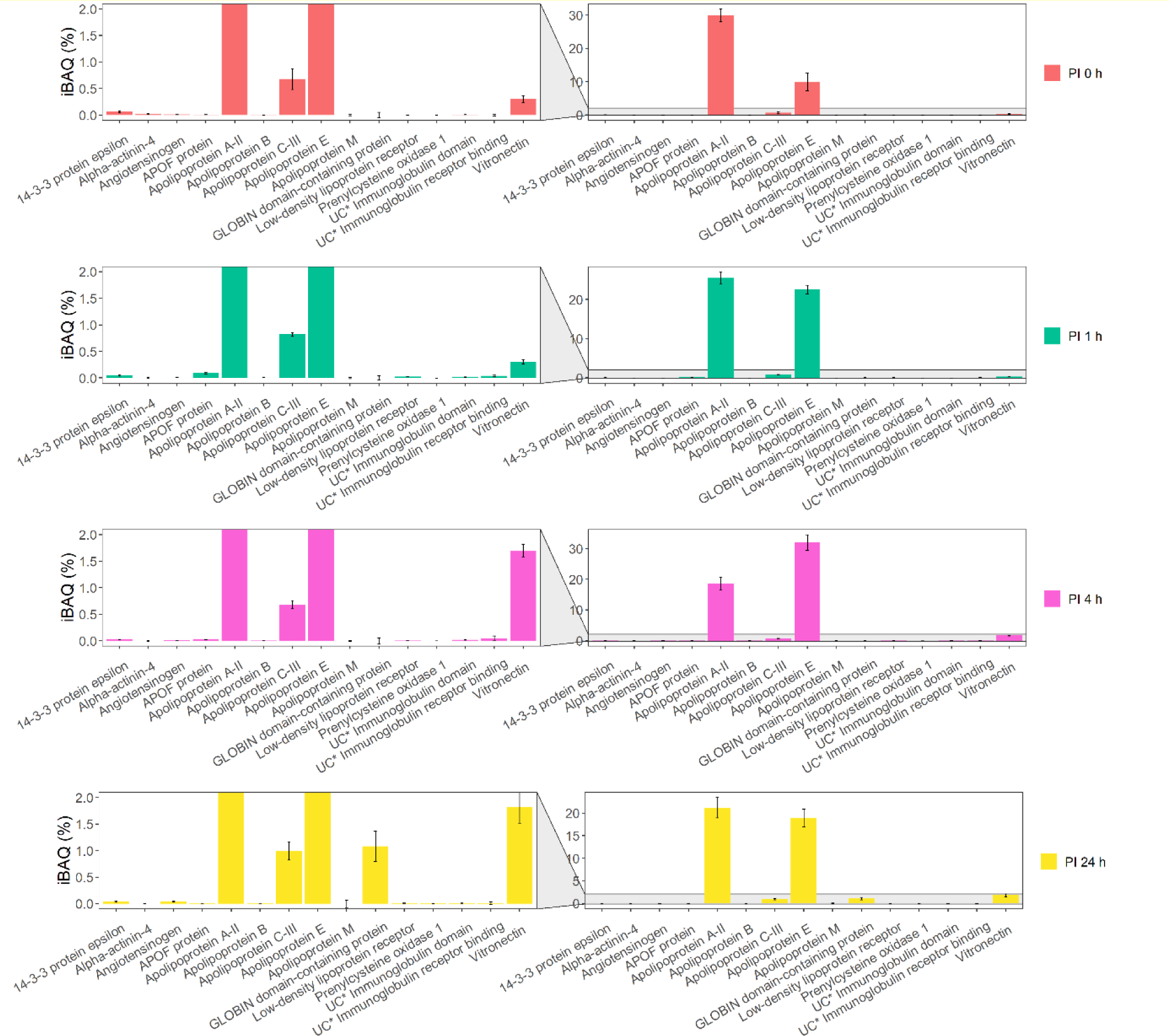
Percentages of the intensity Based Absolute Quantification (iBAQ). Values across the 15 significant proteins identified for the selected PI times. Data from two independent samples analyzed in duplicate (N=2, n=2).

**Table S1.** **Lipid nanoparticles used in the study.** Lipid composition as well as cargo type and encapsulation rate are listed for all LNPs used in the present study. Basic size characterization was conducted using conventional dynamic light scattering (DLS) and nanoparticle tracking analysis (NTA) and mRNA encapsulation was determined using the RiboGreen assay.

|  | **MC3_DMPE**<br>**PEG** | **MC3_DMPE**<br>**PEG-biotin** | **MC3_DMPE**<br>**PEG<sup>+PolyA</sup>** | **MC3_DSPE**<br>**PEG** |
| --- | --- | --- | --- | --- |
| Number of batch used | (n=3) | (n=1) | (n=1) | (n=1) |
| **Composition** |  |  |  |  |
| **Lipids (mol%)** |  |  |  |  |
| Dlin-MC3-DMA | 50 | 50 | 50 | 50 |
| Cholesterol | 38.5 | 38.5 | 38.5 | 38.5 |
| DSPC | 10 | 10 | 10 | 10 |
| DMPE-PEG2000 | 1.5 | 1.494 | 1.5 | - |
| DSPE-PEG2000 | - | - | - | 1.5 |
| DSPE-PEG2000-biotin | - | 0.006 | - | - |
| **mRNA [ratio]** |  |  |  |  |
| eGFP mRNA | 4 | 4 | - | 4 |
| Cy5 eGFP mRNA | 1 | 1 | - | 1 |
| PolyA | - | - | 5 | - |
| **Characterization** |  |  |  |  |
| **Diameter size (nm)** |  |  |  |  |
| DLS (z-avg) | 85 ± 5 | 84 | 101 | 88 |
| Polydispersity Index | 0.047 ± 0.024 | 0.085 | 0.333 | 0.032 |
| NTA | 72.2 ± 0.3 | 88.5 ± 2.2 | 84.1 ± 0.4 | N.D. |
| **mRNA encapsulation** |  |  |  |  |
| Encapsulation (%) | 98 ± 1 | 97 | 99 | 98 |
| Concentration (µg/mL) | 124 ± 11 | 134 | 123 | 139 |
| **Particle concentration** |  |  |  |  |
| NTA (particle/mL) | 7.4 e+12<br>± 3.0 e+12 | 6.5 e+12<br>± 0.4 e+12 | 2.2 e+13<br>± 0.1 e+13 | N.D. |
| **Experiment conducted** |  |  |  |  |
| Imaging, flow cytometry | X |  |  | X |
| Proteomics | X |  |  |  |
| TIRF |  | X |  |  |
| NMR |  |  | X |  |
| SANS | X |  |  | X |
| DLS study in medium | X |  |  |  |

N.D.; not determined

**Table S2.** Absolute size (radii, nm) of LNPs prior to addition of serum as obtained by fitting of SANS data and from DLS

| SANS |  | DLS<sup>a</sup> (Z-average) |
| --- | --- | --- |
|  | **MC3_DMPE PEG** |  |
| Polar radii (nm) | 18.9 ± 0.23 |  |
| Equatorial radii (nm) | 33.3 ± 0.35 | 39.1 ± 0.6 |
|  | **MC3 DSPE PEG** |  |
| Polar radii (nm) | 21.9 ± 0.26 |  |
| Equatorial radii (nm) | 40.0 ± 0.48 | 48.9 ± 0.5 |

<sup>a</sup> DLS measurements assume spherical particles and therefore detect the largest particle radius – in this case the equatorial radius. <sup>[68]</sup>

**Table S3.** FBS-induced changes in the radii and volume of LNPs as obtained by fitting of SANS data.

|  |  |  |
| --- | --- | --- |
|  | **MC3_DMPE PEG** | **MC3 DSPE PEG** |
| Polar radii change (nm) | - 3.47 ± 0.52 | - 4.17 ± 0.46 |
| Equatorial radii change (nm) | + 0.66 ± 0.91 | +2.65 ± 1.12 |
| Volume change (%) | - 32 ± 4.9 | - 30.1 ± 3.9 |

## Notes

### Competing Interest Statement

A. Gallud, M.J. Munson, K. Liu, L. Lindfors, A. Collen and A. Sabirsh are employees of AstraZencea.

